# *Fusarium solani* species complex genomes reveal bases of compartmentalisation and animal pathogenesis

**DOI:** 10.1101/2022.03.22.485422

**Authors:** Daphne Z. Hoh, Hsin-Han Lee, Naohisa Wada, Wei-An Liu, Min R. Lu, Cheng-Kuo Lai, Huei-Mien Ke, Pei-Feng Sun, Sen-Lin Tang, Wen-Hsin Chung, Ying-Lien Chen, Chia-Lin Chung, Isheng Jason Tsai

**Author notes:** Corresponding author: Isheng Jason Tsai.

## Abstract

The *Fusarium solani* species complex (FSSC) comprises fungal pathogens responsible for mortality in a diverse range of animals and plants, but their genome diversity and transcriptome responses in animal pathogenicity remain to be elucidated. We sequenced and compared six chromosome-level FSSC clade 3 genomes of aquatic animal and plant host origins and revealed a spectrum of conservation patterns in chromosomes categorised into three compartments: core, fast-core (FC), and lineage-specific (LS). Each chromosome type varied in structural architectures, with FC and LS chromosomes containing significantly higher proportions of repetitive elements and methylation levels than core chromosomes, with genes exhibiting higher *d*N/*d*S and enriched in functions related to pathogenicity and niche expansion. Mesosynteny were detected between FC chromosomes of *Fusarium* genomes, indicating that these chromosomes were present in a common ancestor that predated FSSC species. These findings provide evidence that genome compartmentalisation was the outcome of multi-speed evolution amongst FSSC chromosomes. We further demonstrated that *F. falciforme* and *F. keratoplasticum* are opportunistic pathogens by inoculating *Pelodiscus sinensis* eggs and identified differentially expressed genes also associated with plant pathogenicity. These included the most upregulated genes encoding the CFEM (Common in Fungal Extracellular Membrane) domain. The study establishes genomic resources and an animal model for fungal pathogens of trans-kingdom hosts.

## Introduction

The filamentous fungi in the genus *Fusarium* are among the most virulent pathogens affecting multi-kingdom hosts (O’Donnell et al., 2016; Zhang et al., 2006), but can also exist as saprophytes. The genus, commonly identified as plant pathogens, has caused devastating losses in the global agricultural industry (Dean et al., 2012) and is recognized as one of the most prevalent clinical pathogens causing superficial and invasive disease in immunocompromised humans (Al-Hatmi et al., 2016; Walther et al., 2017). In recent decades, an increasing number of fusariosis cases associated with various types of animals have been reported worldwide (Brown et al., 2020; Cabañes et al., 1997; Fernando et al., 2015; Hsu et al., 2021; Kano et al., 2002), but research on its virulence beyond plant hosts remains limited (Coleman, 2016; Rep & Kistler, 2010). It is now considered as a serious emerging fungal threat potentially inducing host population loss and extinction (Fisher et al., 2012; O’Donnell et al., 2016).

The most prevalent *Fusarium* pathogens associated with veterinary infection are from the species-rich clade three of *Fusarium solani* species complex (FSSC; O’Donnell et al., 2008; O’Donnell et al., 2016; Schroers et al., 2016; Zhang et al., 2006), which are ubiquitous in the environment (Zhang et al., 2006). Several FSSC species were reported to cause disease in aquatic animals such as grey seals, shrimps, dolphins, and sharks (O’Donnell et al., 2016). Two species from the FSSC clade 3–*F. falciforme* and *F. keratoplasticum*–were the predominant species occurring in diseased sea turtle nests worldwide (Sarmiento-Ramírez et al., 2014), and the latter was also recently reported to infect *Podocnemis unifilis*, an endangered freshwater turtle species (García-Martín et al., 2021). Koch’s postulates were fulfilled for *F. keratoplasticum*, which can cause disease and high mortality rate (83.3 %) in sea turtle *Caretta caretta* eggs (Sarmiento-Ramírez et al., 2010). The disease is now termed sea turtle egg fusariosis (Smyth et al., 2019); it is responsible for low hatching success of eggs from both natural nests (Phillott et al., 2004; Sarmiento-Ramírez et al., 2014) and man-made hatcheries (Hoh et al., 2020; Sidique et al., 2017).

One particular genome characteristic in many pathogenic fungi are that their chromosomes can be differentiated into two compartments: the core chromosome (CC), which contain essential genes required for survival and reproduction, and lineage-specific chromosome (LSC; also known as accessory chromosome), which is repeat-rich and contains enriched genes mostly associated with niche adaptation and pathogenicity (Bertazzoni et al., 2018; Möller & Stukenbrock, 2017). LSC can be dispensable and do not affect fungal growth in several *Fusarium* species such as *F. oxysporum* f. sp. *lycopercisi* (Ma et al., 2010) and *F. vanettenii* (previously *Nectria haematococca*; Wasmann & VanEtten, 1996). The LSC of *F. vanettenii* harbours pea-specific pathogenic genes–the PEP cluster (Han et al., 2001; Temporini & VanEtten, 2002; Wasmann & VanEtten, 1996), and the deletion of the entire LSC reduced the pathogen’s virulence towards pea plants (Coleman, 2016; Wasmann & VanEtten, 1996). It is unclear whether any of these virulence genes are present in other FSSC species, if they play a role in animal infection, and if they are enriched in LSCs.

Here, we show that *F. falciforme* and *F. keratoplasticum* can penetrate eggshells and colonise egg inclusions. We produced six high-quality genome assemblies for species in the FSSC clade 3 and divided the chromosomes into multiple compartments based on different genome features and selection pressures. We developed the first study model for the infection of animal pathogenic *Fusarium* employing the dual RNA-seq sequencing method to identify regulated host and pathogen genes during the infection. Together the results provided a clearer understanding of genomic bases of FSSC species and their virulence mechanisms on animals.

## Results

### *Fusarium falciforme* and *F. keratoplasticum* are opportunistic pathogens of turtle eggs

We analysed the infection scenarios of *F. falciforme* Fu3 and *F. keratoplasticum* Fu6 inoculated in eggs of the animal host Chinese softshell turtle (*Pelodiscus sinensis*). Hyphal growth in both species was observed on the eggshell surface after five days of inoculation (**Figure 1**) with some occasionally growing into the cavity-like structures (**Supplementary Figure 1**). Cryo-sectioning and histological observations of undecalcified eggshell cross-sections revealed the presence of both *Fusarium* species on the outer, within the calcareous, and inner layers of the eggshells (**Figure 1**), confirming that hyphae vertically penetrated the eggshell. Degradations were sometimes observed on the eggshell membrane (**Figure 1**). Attraction assays revealed no significant difference in hyphal growth rates with or without the presence of turtle eggs in each *Fusarium* species (*p* = 0.67 in *F. falciforme* and *p* = 0.86 in *F. keratoplasticum*; **Supplementary Figure 2**) or between species (*p* = 0.86 in control and *p* = 0.86 in treatment). We further examined the symptoms of *F. falciforme* and *F. keratoplasticum* colonisation on eggs at three- and four-days post-inoculation (dpi) (**Supplementary Figure 3**). Mycelial mass was observed growing on the membrane (**Supplementary Figure 3a**) and embryo (**Supplementary Figure 3b and c**). Some inoculated eggs exhibited reduced branching points and disrupted blood capillaries on the microvascular system despite the embryo still being alive during the examination (**Supplementary Figure 3c**). Together, these observations suggested that these two FSSC species are opportunistic animal pathogens and, upon contact, their hyphae can penetrate eggshells via natural openings and subsequently lyse and colonise egg inclusions.

**Figure 1.**
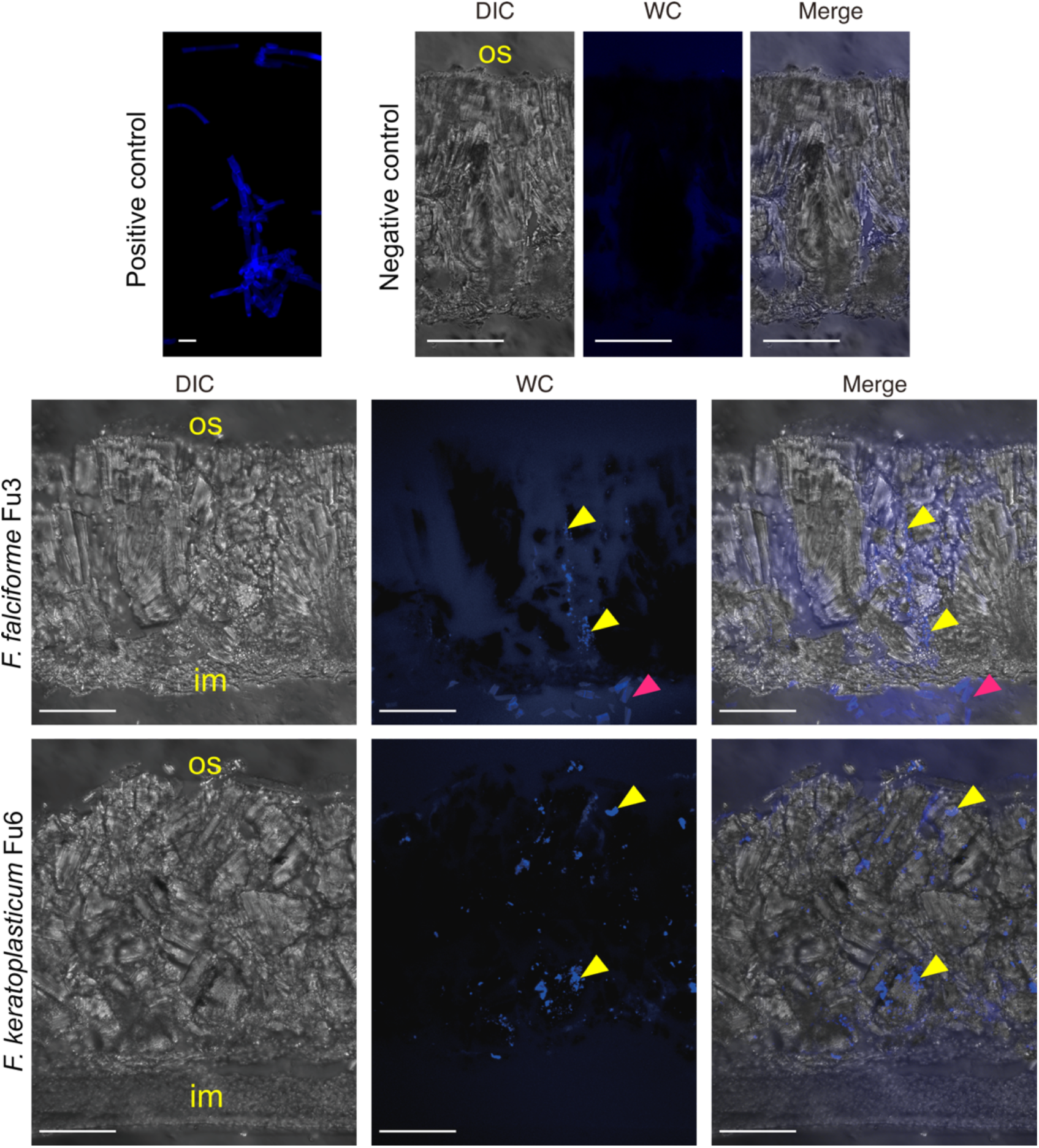
Laser confocal microscopy images of undecalcified cross-section of *Pelodiscus sinensis* eggshell acquired at 5-dpi of *Fusarium falciforme* Fu3 and *F. keratoplasticum* Fu6.

The text ‘os’ and ‘im’ indicate outer eggshell surface and inner eggshell membrane, respectively. Fungal material was stained with Calcofluor White (blue signal). Yellow and pink arrowheads denote fungal signal and degraded eggshell membrane, respectively. DIC = differential interference contrast. WC = white contrast. Scale bar in figure is 50μm except 10μm in positive control.

### Genome characteristics of six sequenced FSSC isolates

We sequenced the genomes of five species within clade 3 of FSSC (**Table 1**). This included *F. falciforme* (Fu3), *F. keratoplasticum* (Fu6 and LHS11), and *Fusarium* sp. FSSC12 (LHS14) isolated from various aquatic animal hosts. Two species of plant host origins–*Fusarium* sp. (Ph1) from orchid and *F. vanettenii* (Fs6) from pea–were also chosen for comparative purposes. The initial assemblies were produced from averaging 121X of Oxford Nanopore reads (sequence N50 = 14–23kb) using the Flye assembler (Kolmogorov et al., 2019) and polished by Illumina reads (**Supplementary Table 1**). The final assemblies were in 14 to 40 pieces with N50 3.2–4.2 Mb (**Table 1**) averaging 56 Mb, which is more contiguated and larger than other representative *Fusarium* genomes (ranging 12–4,197 contigs averaging 47Mb; **Supplementary Table 2**). Interestingly, *F. vanettenii* Fs6 has an assembly of 72.9 Mb, the largest *Fusarium* genome reported to date and larger than the published *F. vanettenii* MPVI 77-13-4 genome of 51.2 Mb (Coleman et al., 2009), suggesting high intraspecies variation. The *F. falciforme* Fu3 reference was amongst the most complete genome, consisting of 14 contigs with six telomere-to-telomere gapless chromosomes (**Supplementary Table 3**), and was on average 17 times more contiguated than the published FSSC genomes (N90: 2.7 Mb vs. 0.4 kb–2.6 Mb, respectively; last accessed date 18^th^ January 2022; **Supplementary Table 4**).

**Table 1.** Genome statistics of six sequenced isolates of *Fusarium solani* species complex.

| Species | *F. falciforme* | *F. keratoplasticum* | *F. keratoplasticum* | *Fusarium* sp. FSSC12 | *Fusarium* sp. | *F. vanettenii* |
| --- | --- | --- | --- | --- | --- | --- |
| Strain ID | Fu3 | Fu6 | LHS11 | LHS14 | Ph1 | Fs6 |
| Genome size (Mb) | 53.6 | 49.5 | 53.5 | 56.3 | 51.5 | 72.9 |
| Number of sequences | 14 | 25 | 26 | 35 | 24 | 40 |
| Largest contig (Mb) | 6.8 | 6.4 | 6.6 | 6.1 | 5.4 | 6.7 |
| N50 Mb (L50) | 3.8 (6) | 4.2 (5) | 3.4 (6) | 3.8 (6) | 3.2 (6) | 3.3 (9) |
| N90 Mb (L90) | 2.7 (12) | 2.2 (11) | 1.5 (13) | 0.9 (14) | 1.1 (17) | 1.3 (23) |
| GC % | 49.4 | 51.3 | 51.4 | 48.6 | 51.1 | 49.4 |
| Number of protein coding genes | 15,137 | 14,935 | 15,051 | 15,937 | 15,498 | 18,862 |
| Protein BUSCO % (fungi) | 99 | 98.4 | 98.4 | 98.3 | 100 | 98.6 |
| Protein BUSCO % (ascomycota) | 98.8 | 98.8 | 98.1 | 98.6 | 99.8 | 98.7 |

We predicted 14,927 to 18,862 protein-encoding genes from these assemblies using the MAKER2 (Holt & Yandell, 2011) pipeline based on evidence using proteomes from closely related fungi and transcriptome reads generated from mycelium. Analysis of Benchmarking Universal Single-Copy Orthologs (BUSCO; Manni et al., 2021) indicated that the proteomes were 98.3 to 100% complete (**Table 1**) and 70–77.1% of these genes were assigned to a protein family or domain (Pfam) via eggNOG-mapper (Cantalapiedra et al., 2021), indicating high completeness of these assemblies. The orthology of the six sequenced assemblies, 17 other *Fusarium* genomes, and an outgroup species *Beauveria bassiana* (**Supplementary Table 2**) was inferred using OrthoFinder (Emms & Kelly, 2019). A total of 24,203 orthogroups (OGs) were inferred, of which 16,154 (66.7%) and 235 (1%) OGs were *Fusarium* and FSSC-specific, respectively. All six genomes contained a predicted average of 420 (2.6% of total gene) and 487 genes encoded for effector and carbohydrate active enzyme, respectively (**Supplementary Table 5 and 6)**. In addition, an average of 44 secondary metabolite biosynthetic gene clusters were detected in the six genomes; of which some included fusarin were associated with plant pathogenicity (**Supplementary Table 7 and 8**; Ma et al., 2013). All these additional gene annotations revealed a similar number of gene features typically comparable to other *Fusarium* genomes (Ma et al., 2013; Niehaus et al., 2016). The repeat proportion of FSSC genomes averaged 7.1% (ranged 4.2–9.4%), except for *F. vanettenii* Fs6, which contained 18.8% repeats (**Supplementary Figure 4a**). The unusually large repetitive DNA content and gene number possibly contribute to the large genome size of *F. vanettenii* Fs6. DNA transposons constituted the largest proportion (2.1 to 6.0%) of FSSC genomes (**Supplementary Figure 4b**), followed by LTR retrotransposons (0.9 to 5.5%). Examination of 5-methylcytosine (5mC) in DNA revealed that methylation levels were similar between coding regions and repetitive elements (average 4.7 vs. 6.5%; Wilcoxon rank-sum test, *p* = 0.2; **Supplementary Table 9**).

Species delimitation within FSSC is challenging because isolate morphologies are almost indistinguishable and nucleotide identities of marker genes are highly similar (O’Donnell et al., 2008). We compared the sequence identity and constructed a maximum likelihood phylogeny based on 40 FSSC species using the commonly employed multi-locus sequence typing (MLST) targeting internal transcribed region (ITS), translation elongation factor (TEF), and RNA polymerase II (RPB2) (**Figure 2a; Supplementary Figure 5 and Table 10**). Even though most of the species were resolved based on the MLST phylogeny, pairwise comparison of sequence identity between some FSSC species was higher than 98%, even between different species, consistent with the limitation of using only a few molecular markers. We further determined the genome average nucleotide identity (ANI) and detected a lower average of 94.5% among the FSSC species comparisons. For instance, *F. falciforme* Fu3 and *F. keratoplasticum* Fu6 had a nucleotide similarity of 98.6% in the MLST sequence but 94.7% in the genome; this better distinction allowed us to resolve FSSC species at the genome level. Ph1 was most closely related to *F. solani* haplotype FSSC5 (MLST and genome ANI: 98.2 and 94.7%). *Fusarium* sp. LHS14 was designated as haplotype FSSC12 and the first genome assembly of this species (**Supplementary Figure 5**). Finally, we constructed a species phylogeny using 2,385 single-copy orthologs, which recapitulated the relationship based on the MLST phylogeny, ANI and found that the species relationships were not grouped by animal or plant hosts (**Figure 2b; Supplementary Figure 6 and Table 2**).

**Figure 2.**
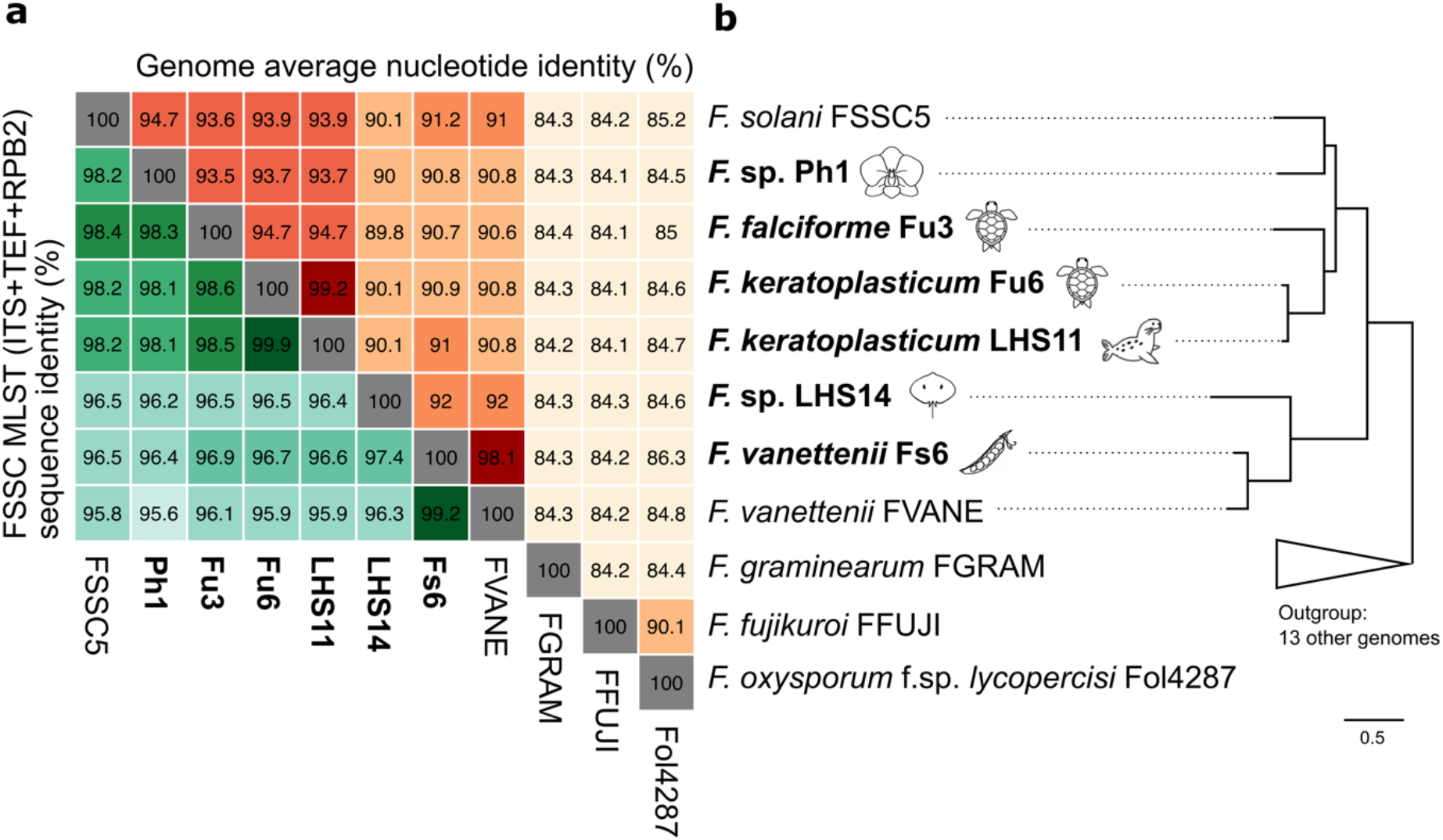
Average nucleotide similarities and species tree of *Fusarium solani* species complex (FSSC). (a) Nucleotide identities determined by multi-locus sequences (ITS+TEF+RPB2) commonly used for FSSC in the lower green-shaded triangular matrix and genome average nucleotide identity in selected *Fusarium* species in the upper orange-shaded triangular matrix. Darker shading indicates higher sequence similarity. (b) A simplified *Fusarium* species tree with outgroup species collapsed. The full phylogeny is in **Supplementary Figure 6** and was constructed using 2,385 single-copy orthogroup sequences. Species name in bold represents the strains sequenced in the current study and source origin (host) represented by icons.

### Evolutionary dynamics of FSSC chromosomes

To investigate whether lineage-specific chromosomes (LSCs) were present in each *Fusarium* species, we assessed the extent of chromosome linkage between species via pairwise single-copy orthologs, proportions of repetitive elements and FSSC-specific genes in each chromosome. In addition to assigning the core and LSC as in previous *Fusarium* studies (Coleman et al., 2009; Ma et al., 2010), FSSC chromosomes were categorised into two compartments, revealing a spectrum of conservation (**Figure 3a**). The majority of chromosomes were part of the linkage group that were shared by all six species; we designated these the core chromosomes (CCs). On the other end of the conservation spectrum were LSCs that contained single-copy orthologues shared by mostly two out of six genomes and a lower proportion of shared genes. Further examination of linkage between FSSC chromosome with three other non-FSSC *Fusarium* species revealed an additional compartment within the FSSC, which had a lower proportion of genes with orthology detected beyond FSSC (**Figure 3b**). This additional compartment comprised three chromosomes (corresponding to chromosome seven, 11, and 12 in five strains and eight, 12 and 13 in *F. vanettenii* Fs6) of FSSC genomes; these were designated as fast-core chromosomes (FCCs). Synteny analysis revealed that the CCs and FCCs were more syntenic than the LSCs between the FSSC species (**Supplementary Figure 7**) and harboured significantly fewer repeats (averaging 5.8, 9.4 vs. 31.3%, respectively; **Figure 4a)**. We designated a total of 9 CCs, 3 FCCs and 2–11 LSCs in each FSSC genome. The CCs and FCCs constituted the majority (67.7–94.7%) of genomes, while the gapless LS chromosomes averaged 1.2 Mb, which is a similar mark up as the *F. oxysporum* genomes (Ma et al., 2010). However, large variations in telomere-to-telomere LSC length were observed in FSSCs, ranging from 0.8 Mb in chromosome 13 of *Fusarium* sp. Ph1 to 3.3 Mb in chromosome 14 of *F. vanettenii* Fs6, demonstrating the characteristic dynamics of non-CCs in *Fusarium* genomes (Fokkens et al., 2018; Zhang et al., 2020).

**Figure 3.**
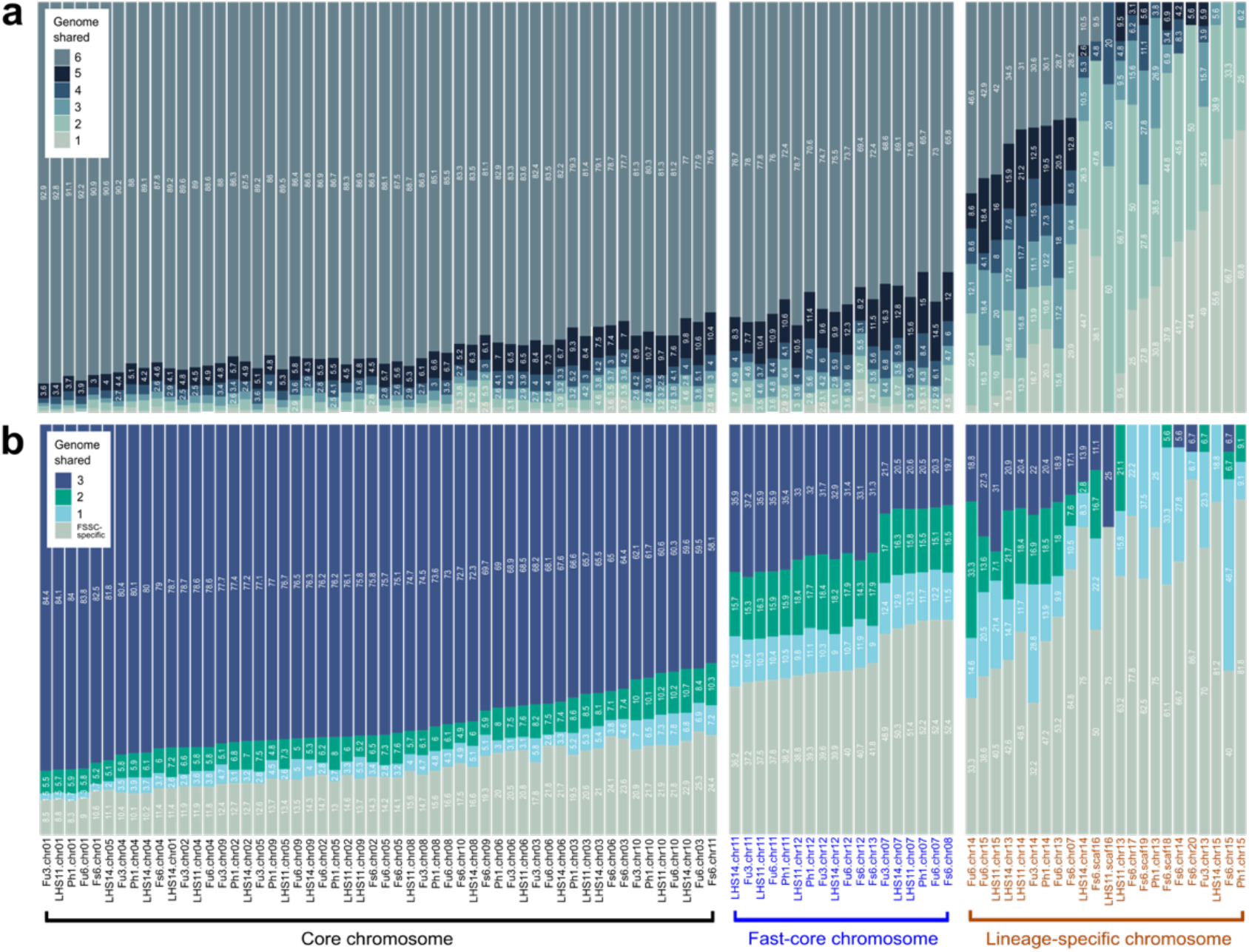
Orthologue sharing amongst *Fusarium* chromosomes. (a) Proportion of one-to-one orthologue shared across each chromosome. (b) Proportion of one-to-one orthologues with three additional *Fusarium* genomes outside of FSSC including *F. oxysporum*, *F. graminearum*, and *F. fujikuroi*.

**Figure 4.**
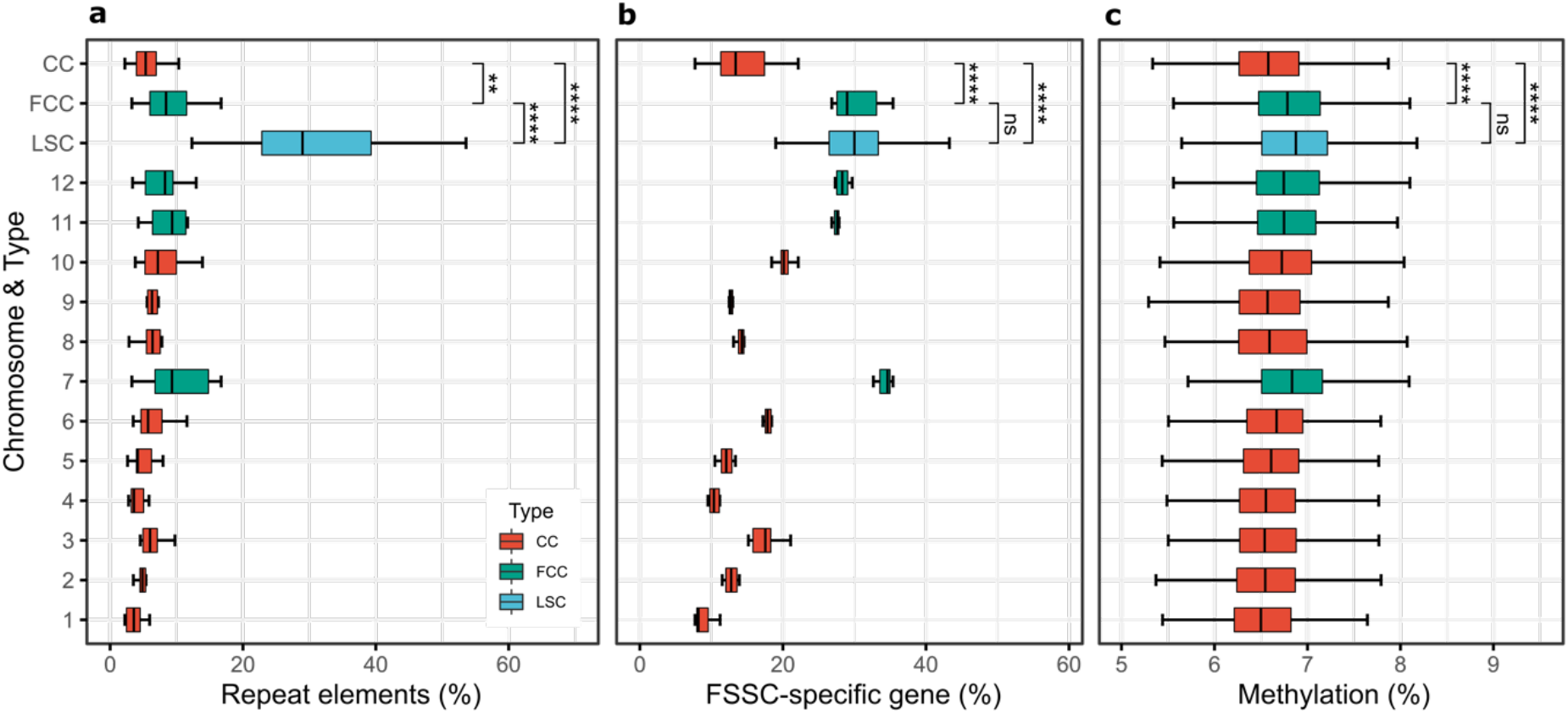
Structural features in *Fusarium solani* species complex (FSSC). Boxplots show the proportions of (a) repeat elements and (b) FSSC-specific gene of each chromosome and chromosome type of the six FSSC genomes. (c) DNA methylation levels of *F. falciforme* Fu3 chromosomes. Outliers lower than 5% and higher than 9% were excluded from the plot. Statistical significance was calculated using Wilcoxon rank-sum test (**: *p* < 0.01; ****: *p* < 0.0001; ns: *p* > 0.05). CC, Core chromosome. FCC, Fast-core chromosome. LSC, Lineage-specific chromosome.

We observed consistent proportions of FSSC-specific genes across corresponding chromosomes of each FSSC genome, with a significantly higher proportion of FSSC-specific genes in FCCs than CCs (**Figure 4b**). The combined mean size of FCCs was approximately 2.8Mb, with FSSC-specific genes ranging from 26.8 to 35.4% of the total gene content (vs. 7.7 to 22.1% in CCs). A lower number of FSSC-specific genes were detected in LSCs compared to CCs (averaging 59 vs. 272 genes), but the former constitutes at least one-fifth of the total gene content (**Supplementary Table 11**). In terms of gene locality, sub-telomeric bias was detected for these FSSC-specific genes, a similar trend as previously reported in *F. graminearum* (Cuomo et al., 2007) and *Aspergillus* genomes (Fedorova et al., 2008), but not in FCCs, where the genes were distributed across the entire chromosome (**Supplementary Figure 8**). Gene Ontology (GO) enrichment analysis in each chromosome type revealed FCCs and LSCs harbours genes that are feasibly linked to pathogenicity and expansion of new niches such as environment or host, compared to CCs which mainly harbours genes for essential biological functions such as growth and development (**Supplementary Information**). FCCs had the highest mean proportion and number of effectors, carbohydrate active enzymes and secondary metabolite biosynthetic genes clusters among the chromosome types, suggesting these genes in FCCs play an important role in pathogenicity processes during host colonisation and infection (**Supplementary Table 12 and 13**). We also observed significantly higher methylation levels in FCCs and LSCs than CCs (**Figure 4c; Supplementary Figure 9**).

To investigate the evolutionary differences of each chromosome type, we estimated the ratio of non-synonymous substitutions to synonymous substitutions per gene (*d*N/*d*S) across single-copy ortholog between *F. falciforme* Fu3 and *F. keratoplasticum* Fu6. Overall, genes located in LSCs had the highest *d*N/*d*S than FCCs, followed by CCs (median ω = 0.24, 0.10, and 0.06, respectively, Wilcoxon rank-sum test, *p* < 0.05 in all comparison pairs; **Figure 5a**), indicating different levels of purifying selections amongst chromosome type. In addition, genes that were translocated (**Supplementary Figure 7**) also comprised of higher *d*N/*d*S (median ω = 0.09, 0.08, and 0.05, in LSCs, FCCs, and CCs, respectively; Wilcoxon rank-sum test, *p* < 0.05 in all comparison pairs; **Supplementary Figure 10)**, suggesting elevated capability of these translocated genes towards adaptation.

**Figure 5.**
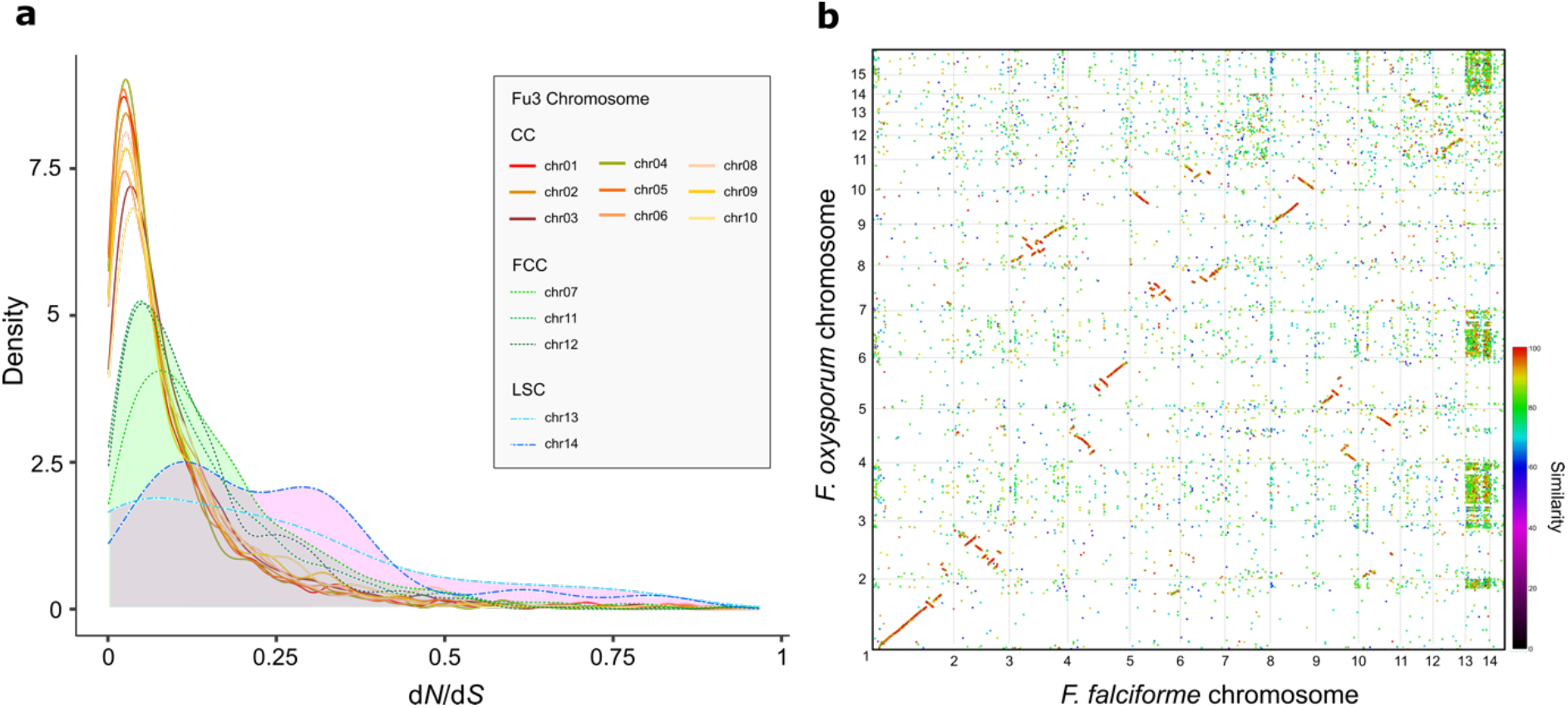
Evolutionary dynamics and origin of fast-core chromosomes. (a) The density of *d*N/*d*S in *F. falciforme* Fu3 chromosomes for each single-copy ortholog gene paired with *F. keratoplasticum* Fu6. (b) Syntenic dotplot produced via PROmer (v3.23; Kurtz et al., 2004) comparing between *F. falciforme* Fu3 and *F. oxysporum* f. sp. *lycopercisi* 4287 genomes. CC = Core chromosome, FCC = Fast-core chromosome, LSC = Lineage-specific chromosome.

Several distinctive differences in chromosome seven (chromosome eight of Fs6) of FSSC genomes were observed when compared to the CCs. For instance, the former was mostly located at the end of FCCs in the conservation spectrum (**Figure 3**); it had the highest proportions of repeat and FSSC-specific genes (**Figure 4a and b**) and higher *d*N/*d*S ratio than other FCCs and CCs (**Figure 5a**). Furthermore, frequent rearrangements were observed in chromosome seven across FSSC genomes (**Supplementary Figure 7b**). Interestingly, intraspecies chromosomal structural variations were observed between *F. keratoplasticum* Fu6 and LHS11 in which translocations of *F. falciforme* Fu3 chromosome seven were observed in chromosome three, thirteen and fourteen of *F. keratoplasticum* Fu6 but dissimilar in LHS11 (**Supplementary Figure 7b**). Hence, the shorter length of chromosome seven of Fu6 suggested that accessory regions were lost and translocated to other chromosomes in the genome. In addition, synteny between *F. falciforme* Fu3 and outgroup *Fusarium* species were assessed to determine the origin of FSSC of FCCs. In the majority of cases, synteny was not detected in all of the FCCs of *F. falciforme* Fu3 (**Supplementary Figure 11**), however, mesosynteny and degraded synteny was found between FCCs of the *F. falciforme* Fu3 and *F. oxysporum* f. sp. *lycopercisi* strain 4287 (Fokkens et al., 2018) genomes (**Figure 5b; Supplementary Figure 12**), indicating a common origin of FCCs in the FSSCs and other *Fusarium* species.

### Transcriptome profiles of FSSC pathogens during egg infection

To better understand how FSSC pathogens infect aquatic animals, we inoculated *F. falciforme* Fu3 and *F. keratoplasticum* Fu6 on *Pelodiscus sinensis* eggs and compared transcriptome-wide gene expression data of both species. Globally, both species adopted similar colonisation and infection strategy at a transcriptome level while contacting eggs (**Figure 6a, Supplementary Figure 13 and 14; Supplementary Information**) and we determined a total of 4,111 (1,823 up- and 2,362 down-regulated) and 4,185 (2,241 up- and 1,870 down-regulated) genes that were differentially expressed (DE; adjusted *p* value < 0.05) in *F. falciforme* Fu3 and *F. keratoplasticum* Fu6 (**Figure 6b**), respectively. When considering these genes regardless of expression level, each pathogen underwent differential responses during egg infection – GO enrichment revealed that *F. keratoplasticum* Fu6 exhibited reactions more related to pathogenicity such as responses to host, stimulus, and toxic substances, cell adhesion, and regulation of immune system whereas *F. falciforme* Fu3 was mostly ribosome-associated processes such as biogenesis, maturation, and transport (**Supplementary Table 14**). We speculated a larger proportion of genes to be differentially expressed in FCCs and LSCs but observed a reverse pattern where the majority of such genes were located in CCs (Chi-square test: Fu3: 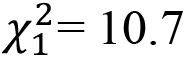, *p* = 0.005; Fu6: 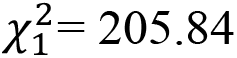, *p* < 0.001). For instance, only 114 and no genes were DE in LSCs of *F. falciforme* Fu3 and *F. keratoplasticum* Fu6, respectively (**Supplementary Figure 15**), indicating genes in FCCs and LSCs were not necessarily enriched in animal pathogenesis.

**Figure 6.**
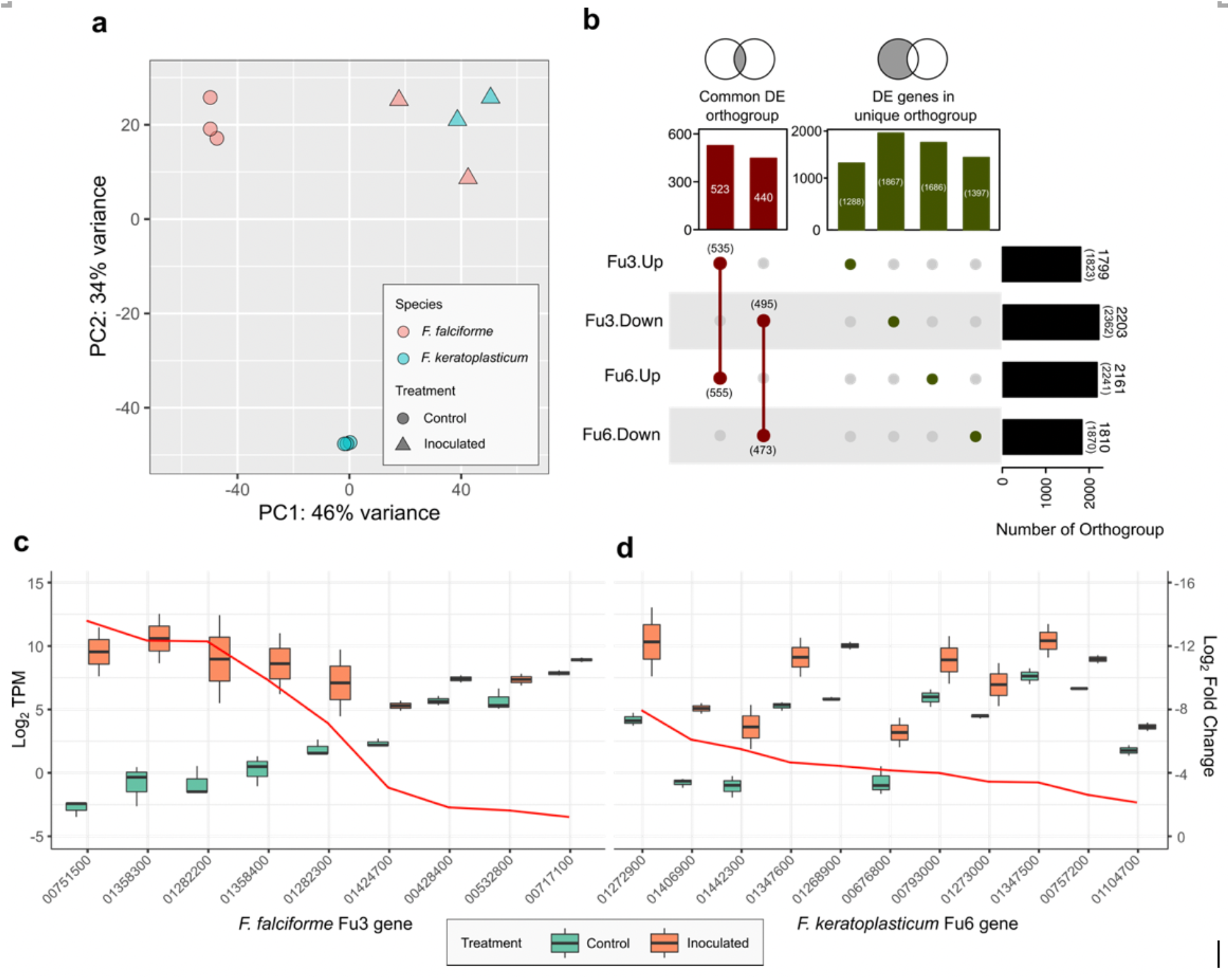
Transcriptomes of *Fusarium solani* species complex pathogens inoculated on animal host *Pelodiscus sinensis* at three- or four-dpi. (a) Principal component analysis (PCA) of gene expression patterns in *F. falciforme* Fu3 and *F. keratoplasticum* Fu6 samples. (b) Number of differentially expressed (DE) orthogroup among the two pathogens. Numbers in the bracket represents number of genes in the orthogroup. (c-d) Expression levels in Log_2_ transcript per million (TPM; left y-axis) and Log_2_ fold change (right y-axis) of genes containing CFEM domain in *F. falciforme* Fu3 (c) and *F. keratoplasticum* Fu6 (d) comparing between control (mycelium grown on PDA) and inoculated samples.

Several genes that have been previously identified to involve in *Fusarium*-plant infection were upregulated in the egg-inoculated samples (**Supplementary Table 15 and 16)**. The majority of these genes were predicted as effectors or contained a signal peptide. Of particular interest, genes annotated to contain CFEM (Common in Fungal Extracellular Membrane) domain, a cysteine-rich protein domain found in diverse phytopathogenic fungal species, comprised some of the most DE genes upregulated in both *F. falciforme* Fu3 and *F. keratoplasticum* Fu6 treatments (11 and 15 genes, respectively; **Figure 6c and d; Supplementary Table 17**). Other examples included the effectors necrosis-inducing proteins (NPP1) and cerato-platanin (CP), which were required for virulence of *F. oxysporum* (Gijzen & Nürnberger, 2006; Liu et al., 2019); ABC membrane and transporter, cytochrome P450 or termed pisatin demethylase (PDA), pelA pectate lyases, whose removal or inactivation reduced the virulence of *F. vanettenii* FSSC11 on pea (Coleman et al., 2011; Rogers et al., 2000; Wasmann & VanEtten, 1996). The results suggested that a similar repertoire of genes were utilised during infection regardless of host types by *F. falciforme* and *F. keratoplasticum*.

In addition, a total of 535 and 555 genes from *F. falciforme* Fu3 and *F. keratoplasticum* Fu6 treatment, respectively belonging 523 OGs were co-upregulated (**Figure 6b**). GO analysis and functional annotations of these genes revealed that both pathogens interacted with the host by positive regulations of their immune systems processes and defence responses (**Supplementary Table 17**). These genes include a TRI12 encoding a major facilitator superfamily protein (MFS_1) involved in the export of mycotoxin trichothecene (Alexander et al., 1999), a nonribosomal peptide synthetase (NPS6), a fungal effector involved in secondary metabolite biosynthesis producing AM-toxin, and PacC, a transcription factor-dependent of pH during pathogenesis (**Figure 7; Supplementary Table 18 and 19**; Caracuel et al., 2003; Lee et al., 2005). In contrast, 440 OGs down-regulated in pathogens during egg infection were involved in transmembrane transports of metal ions, spore development and growth (**Supplementary Table 20**). Furthermore, GO enriched biological processes of species-specific upregulated genes were similar to GO enrichment of all upregulated DE genes in each pathogen species. Upon examination of these upregulated species-specific genes, it was found that 25.7 and 50.4% of the protein domains in genes of *F. falciforme* Fu3 and *F. keratoplasticum* Fu6 respectively, were present in the co-upregulated DE genes of both species (**Supplementary Table 21 and 22**), suggesting similar infective mechanism were adopted by both pathogen species in animal pathogenicity despite diverged gene sequences.

**Figure 7:**
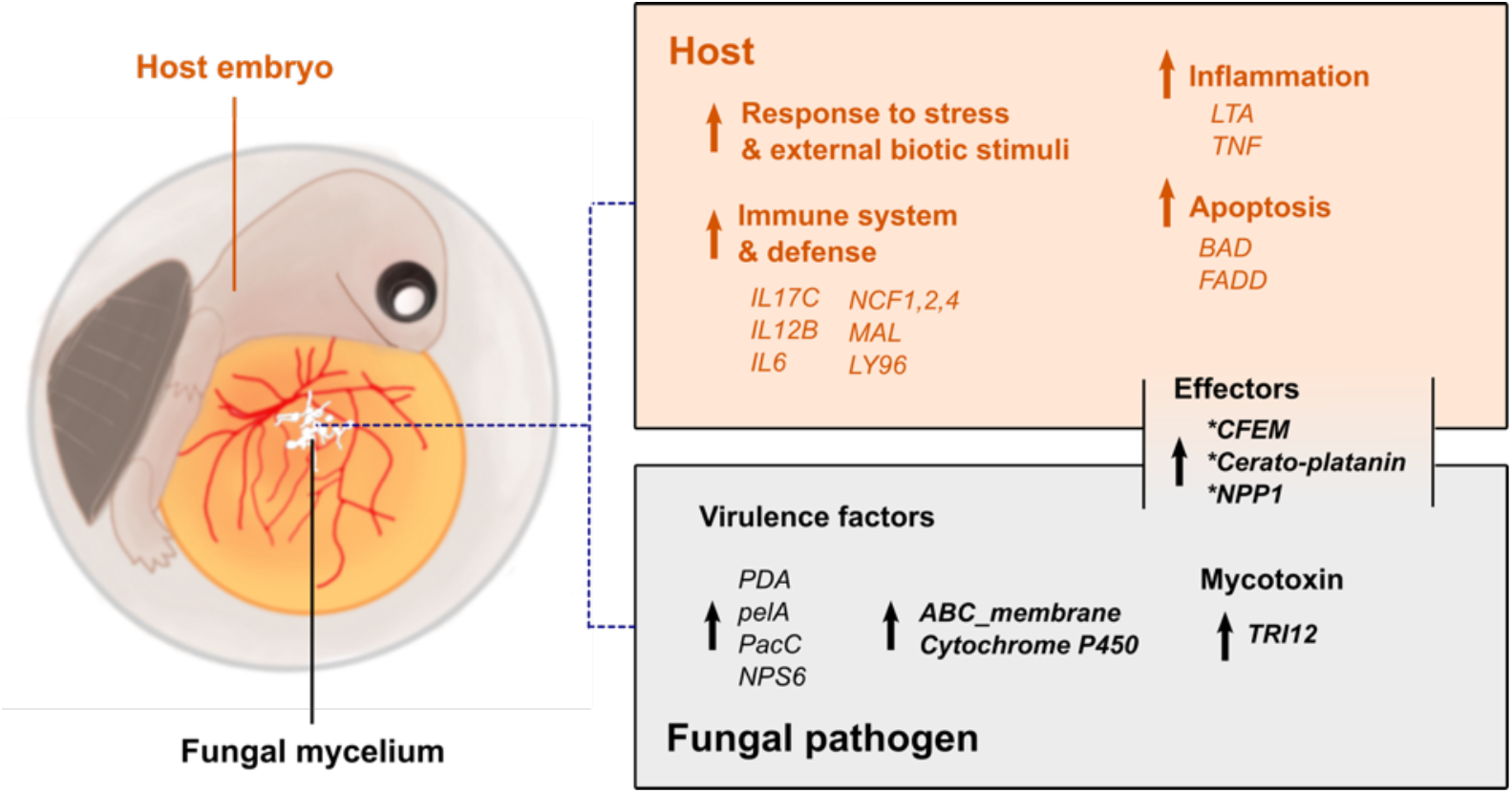
Schematic diagram summarizing genes and functions involved in *Fusarium solani* species complex-*Pelodiscus sinensis* egg infection. The colour-coded text represents turtle host (brown) and pathogens (grey). Up arrow denotes up-regulated genes and enriched functions. Asterisk denotes the presence of signal peptide.

### Gene expression profile of FSSC-infected animal host

Combined with transcriptome data of various development stages in the *P. sinensis* embryos (Wang et al., 2013), the presence (75.0–81.5%) of *F. falciforme*- and *F. keratoplasticum*-inoculated turtle RNA-seq reads in samples allowed us to determine host responses to the two pathogens (**Supplementary Table 23**). A large difference can be seen between the gene expression pattern of the host inoculated by FSSC and the natural developing host embryo on similar egg incubation days (PC1: 95% variance; **Supplementary Figure 16a**), signifying host responded distinctly to FSSC infection. Besides, gene expression in host inoculated with either *F. falciforme* Fu3 and *F. keratoplasticum* Fu6 were highly correlated (*R*^2^ = 0.98, *p* < 0.001; **Supplementary Figure 16b**), suggesting the turtle host responded similarly to FSSC pathogens. In addition to genes that were related to embryo development, other upregulated genes were enriched in biological processes involving host immunity, response, and defence to another organism (**Supplementary Table 24**). Specifically, positive regulations of the immune response towards stress and external biotic stimulus were detected. These regulations included leukocytes activation, cytokines production, apoptotic process, and defence response (**Figure 7**).

## Discussion

Understanding the genetic diversity within FSSC and their ability to infect cross-kingdoms is of fundamental evolutionary interest and essential for the management of emerging wildlife diseases. Here, we produced highly contiguated assemblies for five FSSC species, established the first *Fusarium*-aquatic animal infection model which utilised high-throughput sequencing technology on Chinese softshell turtle (*Pelodiscus sinensis*) egg, and examined gene expression patterns of *F. falciforme* and *F. keratoplasticum* during infection on the animal host. We uncovered diverse evolutionary dynamics of FSSC chromosomes, and these variations were not associated with egg infection. The resources and results of this study allow gained in insights regarding opportunistic infection of FSSC pathogen on animal hosts.

Some FSSC species are model systems for cell biology (Aist & Bayles, 1991), biocatalytic applications (Winkler et al., 2009), and the most extensively studied plant pathosystem that involves *F. vanettenii* on pea (Hadwiger, 2008). Our attraction assays indicated no signs of egg attraction suggesting chance-encounter of FSSC pathogens on eggs, unlike some pathogens that seek hosts actively in the environment (Ruiz et al., 2017; Van Rooij et al., 2015). During plant infection, hyphae of fungal pathogens penetrate host tissue through natural openings such as stomata or lenticels (Ikeda et al., 2019). We observed the initial stage of disease development which include spore germination, hyphal extension and colonisation of *F. falciforme* and *F. keratoplasticum* on turtle eggs, both on the eggshell and internal embryo. While eggshell serves as a protection for the developing embryos, we observed both FSSC pathogens were capable of invading egg inclusions through the eggshell and caused tissue degradation of egg membrane. Previous study had also shown calcium depletion of sea turtle eggshell post-infection by *F. solani* (Phillott et al., 2006), suggesting lytic activity of FSSC pathogens. Nevertheless, the eggshell of *P. sinensis* is thicker than sea turtle eggshell with an expanded calcareous layer (Kusuda et al., 2013), emphasising that egg penetration may have played a more primary role in establishing infection at least for the *P. sinensis* host.

Comparative analyses of the six chromosome-level FSSC assemblies allows the distinction of multiple chromosome compartments based on various structural features. Regions of CCs were highly conserved among the FSSC genomes and macrosynteny were also detected in other non-FSSC species such as *F. oxysporum*, *F. graminearum* and *F. fujikuroi*, indicating these are the CCs across *Fusarium* genera. FCCs harbour several distinctive features compared to CCs which was also described in *F. oxysporum* (Fokkens et al., 2018). We hereby verify multi-compartmentalisation in FSSC genomes in addition to the more commonly discussed “two-speed genome” concept in fungal pathogens (Bertazzoni et al., 2018; Möller & Stukenbrock, 2017; Sperschneider & Dodds, 2022). The mesosynteny detected between FCCs of *F. falciforme* Fu3 and *F. oxysporum* but not in other *Fusarium* species further suggested that such compartmentalisation was ancient and predated the speciation of *Fusarium* species. Additional sequencing of different *Fusarium* strains and species may elucidate the ancestral chromosome feature of FCCs as they displayed differential evolutionary dynamics in each species.

Conserved pathogenicity traits such as effectors in pathogen of different strains might play a role in infecting the same host (Williams et al., 2016). In our egg infection experiment, only a few differences were found between gene expression and overall regulated biological processes of both FSSC pathogens. Interestingly, we detected that plant pathogenicity-related genes were also upregulated in animal infection. These include carbohydrate active enzymes such as cellulase which functions to degrade plant cell wall materials, and pathogenicity genes containing CFEM domain, which were few of the top most expressed genes in both pathogens. CFEM is found in diverse phytopathogenic fungal species with various virulent roles such as appressoria formation in *Pyricularia oryzae* (syn. *Magnaporthe oryzae*; Kou et al., 2017) and root colonisation in *F. oxysporum* (Ling et al., 2015). The most possible explanation for its role in animal infection is biofilm formation, as seen in *Candida albicans* (Pérez et al., 2006) and white blotches surrounding the embryo. Nevertheless, the role of CFEM domain and plant virulence genes in animal infection remained to be elucidated. This combined evidence suggested that FSSC species may adopt a similar repertoire of genes in establishing infection across kingdoms. Interestingly, genes involved in pathogenesis are usually enriched in the LSCs of *Fusarium* species (Coleman, 2016; Dong et al., 2015; H. Yang et al., 2020) but we found no association of differentially expressed genes during egg infection enriched in FCCs and LSCs.

To conclude, the combined results provided new insights into the genomic characteristics of animal-infecting FSSC species and their disease development, particularly on turtle eggs. This research represents the beginning of critical steps to understanding FSSC infection on turtles towards data-informed decision making and management of disease epidemics to reduce disease occurrences in the wild and man-managed settings. Our comparative analyses enabled us to determine multiple compartments in the FSSC genomes indicative of independent evolutionary dynamics across the chromosomes drive for adaptive evolution. Moreover, the reference quality FSSC genomes offer prospects for future research in pathogenesis regardless of the host species.

## Materials and Methods

### Fungal culturing conditions

Six isolates from *Fusarium solani* species complex (FSSC) clade 3 of this study – *Fusarium falciforme* (Fu3), *F. keratoplasticum* (Fu6), *F. keratoplasticum* (LHS11), *Fusarium* sp. haplotype FSSC12 (LHS14), *Fusarium* sp. (Ph1), and *F. vanettenii* (Fs6) underwent the same culture conditions for gDNA and RNA extractions. The isolates were cultured on 1/2 potato dextrose agar (PDA) at 28°C in dark for seven days. *F. falciforme* Fu3 and *F. keratoplasticum* Fu6, which were previously isolated from dead sea turtle eggs, were used to conduct additional pathological experiments that include host attraction assay, animal inoculations and histological observations during disease establishment. For host attraction assay, *F. falciforme* Fu3 and *F. keratoplasticum* Fu6 were cultured on 1.5% water agar at 28°C in dark for five days. *F. falciforme* Fu3 and *F. keratoplasticum* Fu6 cultures prepared for animal inoculations and histological observations experiments were cultured on 1/2 PDA at 28°C in dark for seven days.

### Species identification of isolates

DNA isolation of FSSC mycelium cultures was carried out using ZYMO Quick-DNA Fungal/Bacterial Miniprep Kit (ZYMO Research, Irvine, USA, Cat. #D6005). The identity of these isolates was determined via multi-locus sequence typing (MLST) of ITS rDNA (ITS5), RPB2 (7cF/11aR), and TEF1 (EF1/EF2) regions (O’Donnell et al., 2008), incorporating other FSSC sequences which species identity were determined (**Supplementary Table 10**). PCR conditions for all primers followed Liu et al. (1999) and phylogenetic analysis was performed as described in Hoh et al. (2020).

### Animal sample preparation

Animal experiments were evaluated and approved by the Institutional Animal Care and Use Committee (IACUC) of Academia Sinica, Taiwan under protocol ID 18-05-1214. Freshly laid and fertilized soft-shelled turtle (*Pelodiscus sinensis*) eggs were purchased from a local farm in Pingtung, Taiwan. The eggs were embedded in styrene foam to avoid movement-induced mortality during transportation. The top of the egg surface was marked with a pencil to ensure eggs were not rotated during the following process: egg surface was cleaned with a brush to remove dirt and surface-sterilized by immersing in 75% EtOH for 1 min, 1% bleach + Tween 20 solution for 1 min, and autoclaved distilled water for 1 min. Egg surface was wiped dry using clean tissue paper and half-buried in sterilized and moist vermiculite (1g water/1g vermiculite) in a plastic container covered with cling film. Each container contained ten eggs and was incubated in a climate chamber (Panasonic MLR-352H, Gunma, Japan) at 28°C and 60% relative humidity in the dark until the 30^th^ day. On day 30, the eggs were observed using the candling technique to check for embryo viability. These embryos were estimated to be developed to stage TK21 to 22 (Tokita & Kuratani, 2001). Dead eggs were removed and alive eggs were kept for the following experiments which included pathogen inoculation, host attraction assay and eggshell observations. Any alive embryos remaining after all experiments were humanely euthanized following the standardized procedures approved by IACUC.

### Attraction assays

To determine if the pathogens *F. falciforme* Fu3 and *F. keratoplasticum* Fu6 can be attracted to the egg host, an attraction assay experiment was performed in glass tubes (10cm tube length and 2cm diameter) placed horizontally and filled with approximately 15mL of 1.5% water agar. An approximately 0.5cm^3^ mycelial block was cut from the margin of a 1.5% water agar and carefully transferred to the end of the glass tube without touching the tube surface and bottom agar. At the opening of the glass tube, the egg was fixed using cling film and rubber band in the experimental group while a stopper was used in the control group (**Supplementary Figure 2a**). The horizontally placed tubes were incubated at 28°C in dark and hyphal growth was recorded every day for nine days. The experiment was repeated twice with a total of 38 samples (tubes) assessed.

### Histology during initial disease establishment

*P. sinensis* eggs were inoculated by placing an approximately 1cm^3^ mycelial block (cut from the margin of the colony) on the shell surface and incubated at 28°C in the dark. After five days, the mycelial block was removed and eggshell fragments surrounding the mycelial block were cut and collected for scanning electron microscopy (SEM) and laser scanning confocal microscopy (LSCM) observations. For SEM, eggshell fragments were first fixed with 4% paraformaldehyde and 2.5% glutaraldehyde in 1x PBS buffer for 1 hour at 4°C. Samples were then washed with 1x PBS buffer thrice for 10 min each, followed by second fixation using 1% osmium tetroxide-buffered solution for 1 hour in the dark at room temperature. Fixed samples were washed again as previously described and went through a serial dehydration step using EtOH at 30, 50, 70, 80, 90, 95, and 100% concentration for 10 minutes at each step. Dehydration using 100% EtOH was repeated twice and samples were then dried in Pelco CPD#2400 CO_2_ critical point dryer (Ted Pella Inc., Redding, USA). Finally, samples were coated with a layer of gold with Sputter Coater 108 auto (Cressington Scientific Instruments, Watford, UK) and examined using Quanta 200 ESEM (FEI Company, Hilsboro, USA). For LSCM, eggshells fragments were placed into warm 1.5% agarose and waited until solidified. Each solidified sample embedded in agarose was transferred to a 50mL Falcon tube and fixed with 10% formalin overnight at room temperature. Fixed samples were washed with PBS buffer thrice and then kept in PBS buffer for storage at 4°C until further processing. Sample embedding and undecalcified sectioning procedures followed Wada et al. (2016). Each section was cut into 8μm thickness using an adhesive film and a tungsten carbide blade (SL-T25, Section-Lab Co. Ltd, Japan) on Leica CM3050 cryostat (Leica Microsystems, Nussloch, Germany). The sections on the adhesive film were directly stained with Calcofluor White Stain (Sigma-Aldrich, #18909) for a few minutes, washed with water to remove excess stain, and mounted with ProLong™ Gold Antifade Mountant (Thermo Fisher Scientific, Waltham, MA, USA, #P36930), followed by removing excess mounting solution and covered with a coverslip. Slides were kept at 4°C in the dark until LSM observation. Samples were examined using LSM880 (ZEISS, Germany) and visualized with Zen2.3 software black edition (ZEISS, Germany). Signals of Calcofluor White were excited with 405nm laser and detected in range 410–523nm. DIC (differential interference contrast) images were also acquisted. All image acquisitions were scanned with a 40x objective lens in z-stack mode (0.391μm). The z-stack images were prepared with a maximum intensity projection through Zen2.3 software lite edition (ZEISS, Germany).

### Nucleic acids isolation, genome and transcriptome sequencing

Six isolates from FSSC clade 3 were chosen for genome sequencing and comparative analyses. Fresh mycelium from the cultures was scraped off from the culture media for gDNA and RNA isolation. Genomic DNA was extracted following protocols designed for high-molecular-weight gDNA sequencing (Mayjonade et al., 2016) and size selection and purification of isolated gDNA was performed following (Chouikh et al., 1979). The integrity of gDNA was evaluated using Fragment Analyzer 5200 (Advanced Analytic Technologies, Inc., Ankeny, USA) and the fragment lengths were determined using PROsize 2 software (Advanced Analytic Technologies, Inc., Ankeny, USA). Genomes were sequenced using both Illumina and Oxford Nanopore platforms. Detailed information such as sequencing platforms, library kits used and sequence accession number for each sample can be found in **Supplementary Table 25**. The summary of DNA sequencing data is shown in **Supplementary Table 1**. For gene model prediction and annotation, RNA of the six isolates were extracted following the TRIzol reagent protocol (Thermo Fisher Scientific, Waltham, MA, USA, Cat. #15596026). The integrity of the isolated RNA was checked using 1.5% agarose gel and quantity was determined using Invitrogen Qubit® 4 Fluorometer (Thermo Fisher Scientific, Waltham, MA, USA) before library preparation and sequencing (**Supplementary Table 25**).

### Genome assembly and annotations

Guppy (v3.2.4 and v4.0.11; Oxford Nanopore Technology) or albacore (for 1D^2^ reads; v2.2.7; Oxford Nanopore Technology) were used to perform basecalling of Nanopore raw signals (see **Supplementary Table 1**) and assembled using Flye (v2.5; Kolmogorov et al., 2019) and further polish and correct the assembled sequences with consensus mapped reads from Illumina using Pilon (v1.22; Walker et al., 2014). Haplotypes were collapsed with HaploMerger2 (ver. 20180603; Huang et al., 2017). Annotation of repeat elements of the genomes was performed following Berriman et al. (2018) except described otherwise in the following: a consensus repeat library was created using repeat elements identified via RepeatModeler (v.open-1.0.8) and TransposonPSI (release 08222010; http://transposonpsi.sourceforge.net/) and merged using usearch (v8.1.1861; Edgar, 2010). RepeatMasker (v.open-4.0.7; option -s; https://www.repeatmasker.org) was used to mask the predicted repeat regions in each genome. Telomeric repeats of each scaffold were determined using Tandem Repeat Finder (v4.09; default parameters; Benson, 1999). Enriched hexamers were identified (TTAGGG) manually using Python script on the terminal 5 kb regions of each scaffold to confirm the presence of telomeres. Genes of the assemblies were predicted using AUGUSTUS (v3.3.3; Hoff & Stanke, 2018) and trained with BRAKER2 (option fungi and softmasked; v2.1.4; Brůna et al., 2021). MAKER2 (v2.31.9; Holt & Yandell, 2011) was then used to combine evidence from the assembled transcripts, reference proteomes, and transcript evidence obtained from RNA-seq of the mycelium to produce a final gene annotation set. Completeness of each genome was accessed by BUSCO (v5.2.2; Manni et al., 2021; **Table 1**). Functional annotations of the amino acid sequences were carried out using eggNOG-mapper v2 (http://eggnog-mapper.embl.de; default parameters; Cantalapiedra et al., 2021) on eggNOG v5 database (Huerta-Cepas et al., 2019). Protein domains and families were identified using pfam_scan.pl (v1.6; http://ftp.ebi.ac.uk/pub/databases/Pfam/Tools/) on Pfam database (release 34.0; Mistry et al., 2021). Additional annotations were carried out as follows: carbohydrates active enzymes were determined using dbCAN2 (v2.0.11; Zhang et al., 2018); secondary metabolites detection via antiSMASH (v6.0; Blin et al., 2021); fungal effectors were predicted using EffectorP (v3.0; Sperschneider & Dodds, 2022) on amino acid sequences which passed the signal peptide prediction via signalP (v6.0; Teufel et al., 2021); and genes related to pathogen-host interaction were determined via PHI-base (v4.11; Urban et al., 2020).

### Comparative genomic analyses

Orthology was assigned using OrthoFinder (v2.3.8; Emms & Kelly, 2019) with proteomes of six species from this study, 17 other *Fusarium* published assemblies from and an outgroup species *Beauveria bassiana* (**Supplementary Table 2**). A total of 2,385 single-copy orthogroups were determined from protomes of 24 species and were aligned using MAFFT (v7.487; option -maxiterate 1000; Katoh & Standley, 2013). The alignments from each of the orthogroup were sent to RAxML-NG for maximum likelihood phylogeny inference (v0.9.0; option --model LG+I+F+G4, --bs-trees 100; Kozlov et al., 2019). The maximum likelihood trees and bootstraps-supported replicates generated from the previous step were combined for the construction of consensus species tree using ASTRAL-III (v5.7.7; option -r 100; Zhang et al., 2018). Lastly, a maximum likelihood phylogeny from concatenated amino acid alignments of single-copy orthogroup was built using RAxML-NG (v0.9.0; option --model LG+I+F+G4, --bs-trees 100; Kozlov et al., 2019) with 100 bootstrap replicates. Codon alignments of one-to-one orthologs between *F. falciforme* Fu3 and *F. keratoplasticum* Fu6 were produced using TranslatorX (v1.1; Abascal et al., 2010). Ratio of *d*N/*d*S were calculated for each ortholog alignment using PAML^2^ of CODEML program (v4.9e; option runmode=−2, seqtype=1, CondonFreq=3, and fix_omega=0; Yang, 2007)

### Core, fast-core and lineage-specific chromosomes assignment

Scaffold sequences shorter than 450kb were not included in this assignment. Lineage-specific chromosomes (LSCs) were assigned based on: (1) reduced number or no synteny regions linked by single-copy orthologous genes between chromosomes of the six FSSC species (**Supplementary Figure 7**), (2) higher repeat proportion (**Figure 3a**), and (3) lower GC content compared to the core chromosomes (**Supplementary Table 11**). The designated LSCs include chromosome 13, 14 of Fu3, chromosome 13, 14, 15 of Fu6 and Ph1, chromosome 13, 14, 15 and 16 of LHS11, chromosome 13 to 19 of LHS14 and chromosome 7, 14 to 23 of Fs6. Chromosomal syntenic regions were determined by linking 9,101 single-copy orthologous genes shared among the FSSC genomes. For this analysis, OGs were determined via OrthoFinder (v2.3.8; Emms & Kelly, 2019) using 11 *Fusarium* genomes (six genome assemblies created from this study, *F. vanettenii* 77-13-4, *F. solani* FSSC5, *F. oxysporum* 4287, *F. graminearum* Ph-1, and *F. fujikuroi* IMI58289). Fast-core chromosomes were determined by comparing proportion of FSSC-specific gene in all FSSC chromosomes (**Figure 3b**). FSSC-specific gene were referred to as orthologous genes not found in *F. fujikuroi* IMI58289, *F. graminearum* PH-1, and *F. oxysporum* f. sp. *lycopercisi* 4287 genomes (Cuomo et al., 2007; Ma et al., 2010; Wiemann et al., 2013).

### Methylation analysis

To examine 5-methylcytosine (5mC) in DNA sequences, Nanopore raw FAST5 files of *F. falciforme* Fu3, *F. keratoplasticum* LHS11, *Fusarium* sp. Ph1, and *F. vanettenii* Fs6 were used to run Megalodon (v2.3.3; Oxford Nanopore Technologies) and Guppy (v5.0.11; Oxford Nanopore Technologies) using the default parameters. To determine the methylation level of each CpG site, we calculated the ratio of methylated reads including both strands.

### Animal inoculation with FSSC for transcriptome profiling

During candling observation, the location of the embryo and directly opposite to the embryo (upper pole) were marked. The observed and alive eggs were half-buried on fresh sterilized and moist vermiculite with embryos placed at the lower pole (buried). A tiny hole was carefully made on top of the eggshell and this created a small air space under the hole and above the egg content. Spore suspension of the cultures was prepared by washing the mycelium with 1x PBS solution and filtered through 40μm cell strainer. Haemocytometer-estimated 10^7^ spores/mL suspension was injected into the egg’s air space through the tiny hole without poking the egg content. The inoculated eggs were incubated at 28°C and 60% relative humidity in the dark for another three to four days. The mycelium of *F. falciforme* Fu3 and *F. keratoplasticum* Fu6 collected from 4-day-old colony grown on 1/2 PDA were used as the control. After 3 to 4 dpi, eggs were made broken to check the embryo’s vitality (**Supplementary Table 26**). Samples collected for RNA isolation are of two types depending on the fungal growth during collection: (1) the presence of mycelium mass as white blotches on the egg content (such as embryo and yolk) and (2) cotton-wool-like growth on the opaque eggshell membrane (**Supplementary Figure 3**). Since this experiment focused on the pathogen, we tried to collect only the white substances and the shell membrane with obvious fungal growth. Each sample type was considered one sample. Samples were collected using sterilized forceps and kept immediately in Trizol reagent (Thermo Fisher Scientific, Waltham, MA, USA, Cat. #15596026) and –80°C until further processing.

### Total RNA isolation, library preparation and sequencing

RNA isolation of the animal inoculation experiment was carried out according to the TRIzol reagent protocol. At the cell lysis step, each sample in 2mL screw-cap tube was mixed with six to eight 0.8mm stainless steel beads, flash-freezing the tubes in liquid nitrogen, and homogenized using PowerLyzer24 homogenizer (MoBio Laboratories, Carlsbad, USA, Cat. #13155) set to 3000 x g for 20 sec and repeated at least twice to ensure the sample was homogenized. Isolated RNA was quantified using Invitrogen Qubit® 4 Fluorometer (Thermo Fisher Scientific, Waltham, USA) and checked for integrity in 1.5% agarose gel. Samples with adequate concentration and show no degradation was chosen for RNA sequencing. In total, 20 samples were sequenced, which include three *F. falciforme* Fu3 and three *F. keratoplasticum* Fu6 positive controls (mycelium from PDA media), seven *F. falciforme* Fu3 and seven *F. keratoplasticum* Fu6 infected samples (**Supplementary Table 26**). Paired-end library was prepared for the RNA samples using NED Next® Ultra™ RNA Library Prep Kit and sequenced on Illumina NovaSeq 6000 instrument with 150 bp paired-end reads.

### Analysis of RNA-seq reads

Raw RNA reads were trimmed to remove adaptor sequences and poor-quality reads using fastp (option −l 30; v0.20.1; Chen et al., 2018). Trimmed reads of each sample were mapped to their respective genome (*F. falciforme* Fu3 or *F. keratoplasticum* Fu6) according to the inoculation treatment and the host *P. sinensis* genome (GCF_000230535.1_PelSin_1.0; Wang et al., 2013) using STAR (v2.7.7a; Dobin et al., 2013). To ensure RNA reads which mapped on the *Fusarium* genomes were not from the host, reads mapped onto both *Fusarium* and host genome and had lower CIGAR scores in *Fusarium* were excluded from further analyses. Raw read count of each gene was calculated using featureCounts (-p -s 2; v2.0.1; Liao et al., 2014). Differentially expressed genes (DEGs) between infected and control samples in *F. falciforme* Fu3 and *F. keratoplasticum* Fu6 were inferred using DESeq2 (*padj* < 0.05; v1.24; Love et al., 2014). Functional enrichment of the DEGs was identified using the ‘topGO’ (v2.36.0; Alexa & Rahnenführer, 2019) package. We used the same pipeline as described for the analysis of the host’s RNA reads to determine the DEGs of the infected host using the *P. sinensis* genome (Wang et al., 2013) as reference. For comparison, the RNA-seq dataset of *P. sinensis* embryos from different embryo developmental stages (TK19 and TK23 defined by Tokita & Kuratani, 2001) generated by Wang et al. (2013) was chosen as the control because our samples were collected between TK21 to 22. Of those, 12,419 genes were differentially expressed (adjusted *p* value < 0.05), with 5,815 up- and 6,604 down-regulated genes during the infection experiment.

### Data availability

All sequences generated from this study were deposited on NCBI under BioProject PRJNA782245 and accession number of gene sequences from MLST of isolates can be found in **Supplementary Table 10**.

## Supplemental materials

Supplementary Text

Supplementary Figure 1 to 16

Supplementary Table 1 to 26

## Authors contributions

IJT conceived and led the study; DZH conducted the experiments; DZH, WAL, HMK, and PFS isolated samples’ nucleic acids; DZH, NW and SLT conducted the laser confocal microscopy experiments; HHL and IJT performed assemblies and annotations of FSSC genomes; DZH and IJT carried out comparative genomic analyses; HHL carried out the methylation analyses; CKL carried out the *d*N/*d*S analyses; DZH designed and conducted the inoculation experiments; DZH and MRL analysed the RNA-seq data; WHC and YLC provided FSSC strains. DZH and IJT wrote the manuscript with input from CLC and others. All authors read and approved the final manuscript.

## Acknowledgements

We thank the Department of Fisheries Malaysia, World Wide Fund for Nature Malaysia for sampling permission in Padang Kemunting Turtle Hatchery, Melaka, and Maximus Cheing YC for collecting Fu3 and Fu6 strains. We are grateful to Dr Stéphane Hacquard, Dr Francis Martin, and the ‘1000 Fungal Genomes–Deep Sequencing of Ecologically-relevant Dikarya’ project for access to unpublished genome data. The genome sequence data were produced by the US Department of Energy Joint Genome Institute in collaboration with the user community. We thank the National Center for High-performance Computing (NCHC) for providing computational and storage resources. We thank High Throughput Genomics Core, Biodiversity Research Center Academia Sinica for the sequencing service; Division of Core Facilities Imaging, Institute of Cellular and Organismic Biology, Academia Sinica for SEM sample preparation service; Dr. Po-Yang Chen and Division of Electron Microscope, Cell Biology Core Lab, Institute of Plant and Microbial Biology, Academia Sinica for assistance in using SEM; Dr. Yoko Nozawa for providing histological materials and Neuro-imaging Core, Neuroscience Program of Academia Sinica for LSM imaging service. This work was supported by AS-CDA-107-L01 to IJT. DZH, HHL and PFS are supported by the doctorate fellowship of the Taiwan International Graduate Program, Academia Sinica of Taiwan.

