## Supplementary Info for "*Fusarium solani* species complex genomes reveal bases of compartmentalisation and animal pathogenesis"

This file includes:

1. Supplementary Text: Extended Results
2. Supplementary Figure 1 to 16

Other supplementary materials for this manuscript includes:

- Supplementary Table 1 to 26

### Supplementary Text

#### Extended Results

##### Fast-core and lineage-specific chromosomes are enriched in genes associated with pathogenicity and niche adaptation.

While we expected different structural characteristics and selection pressures on each chromosome type, we asked whether these chromosome types have genes which are distinct in terms of biological functions. We first compared the mean proportion and number of effectors, carbohydrate active enzymes (CAZymes) and secondary metabolite biosynthetic genes clusters among the chromosome types and determined FCCs had the highest among all comparisons (**Supplementary Table 12 and 13**). The findings suggested FCCs were more likely to involve in pathogenicity processes during host colonization and infection. Annotation via Cluster of Orthologous Groups (COG) revealed the CCs had higher proportion of genes related to biological functions associated to RNA processing and modification, nucleotide metabolism and transport, and translation, compared to FCCs and LSCs. Most of the genes in FCCs had COG category associating with metabolism and transport of carbohydrates and amino acids, energy production and conversion, secondary structure, defence mechanisms and biogenesis of cell wall, cell membrane and envelop. Lastly, LSCs had most genes in COG category related to chromatin structure and dynamics, cell cycle control and mitosis, replication and repair, and transport and metabolism of inorganic ions.

We determined significant overrepresentation of genes through Gene Ontology (GO) analysis in each chromosome type and found similar pattern as in COG analysis. CCs contained enriched genes mainly involves in the core biological processes such as regulation, primary metabolism and biosynthesis of organic macromolecules (i.e. proteins and nucleic acids). FCCs had genes associated with processes such as hyphae and mycelium growth, cell wall biogenesis, response to host defences and stress, regulation of immune system processes, detoxification, secondary metabolites (i.e. toxin) biosynthesis, and metabolic process of carbohydrates, amino acids and ions. Finally, enriched genes in LSCs involve in regulation of developmental process such as cell differentiation, cell wall biogenesis, cell morphogenesis, and asexual sporulation. LSCs also have genes involve in processes which response to external stimulus and host, regulation of immune system processes, mycotoxin biosynthesis, and transmembrane transports of various kinds of materials which include ions, cation, and organic acids. In summary, FCCs and LSCs harbours genes which are feasibly linked to pathogenicity and expansion of new niche such as environment and new host, compared to CCs which mainly harbours genes associated to essential cellular functions.

**Gene expression pattern of FSSC during animal infection**

Principal component analysis (PCA) shows that expression pattern of single copy orthologs among *F. falciforme* Fu3 and *F. keratoplasticum* Fu6 pathogens were separated by treatment types (**Figure 6a**; all samples in **Supplementary Figure 13)**, indicating both pathogen species responded distinctly after contacting an animal host compared to culture media. Significant high correlation was found in expression of single copy orthologs in inoculated samples (**Supplementary Figure 14a**; $Log_2$ TPM, $R^2 = 0.79$, $p < 2.2e^{-16}$) which was slightly higher compared to the control samples (**Supplementary Figure 14b**; $Log_2$ TPM, $R^2 = 0.65$, $p < 2.2e^{-16}$), suggesting both species adopted similar colonization and infection strategy while contacting animal host. a total of 1288 and 1867 genes were up-and down-regulated in *F. falciforme* Fu3 treatment, respectively. In *F. keratoplasticum* Fu6 treatment, the up- and down-regulated unique genes were 1686 and 1397, respectively. GO enriched biological processes of upregulated unique genes in each pathogen species were similar as GO enrichment of all upregulated DEGs, in which *F. keratoplasticum* Fu6 displayed a notably pathogenesis-related processes.

**Plant associated pathogenicity genes were upregulated during animal infection.**

We detected several plant pathogenicity associated genes were upregulated during egg infection in both pathogen treatments, these genes encode for cell wall degrading enzymes such as cellulase, chitinase and pectinase. Furthermore, we detected numerous genes containing protein domains which were specifically verified in *F. vanettenii* (FSSC 11) to be associated with pea virulence (Coleman, 2016), were differentially upregulated during animal infection in both pathogen treatments. These protein domains include virulence factors on pea host, which suggest these FSSC virulence genes were not pea plant specific. All these results suggest FSSC pathogens utilize a common infection process regardless of host types from different kingdoms.

#### Supplementary Figure

**Supplementary Figure 1.** Scanning electron microscopy images of Chinese soft-shelled turtle *Pelodiscus sinensis* eggshell inoculated with *F. falciforme* Fu3 or *F. keratoplasticum* Fu6 on five-dpi, showing fungal hyphae spreading on the eggshell surface. Arrowhead indicates hyphae growing into cavity-like structure.

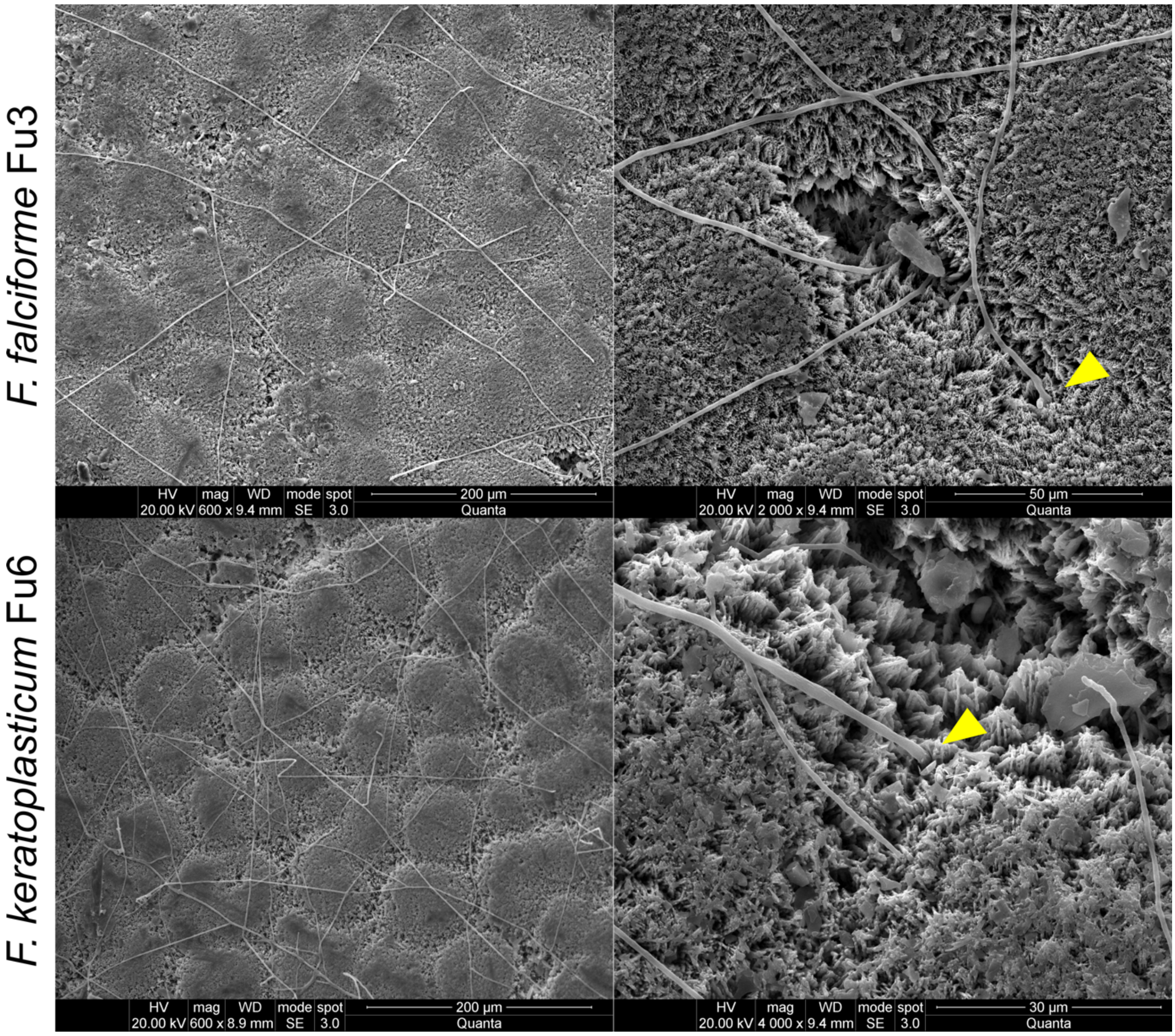

**Supplementary Figure 2.** Host attraction assay. (a) Experimental setup for control treatment using stopper (left tube) and experimental treatment using *P. sinensis* egg (right tube) placed horizontally upon beginning of experiment. (b) hyphae growth rate of all treatments had no significant difference tested using Wilcoxon-test (ns: $p > 0.05$) in each group comparisons.

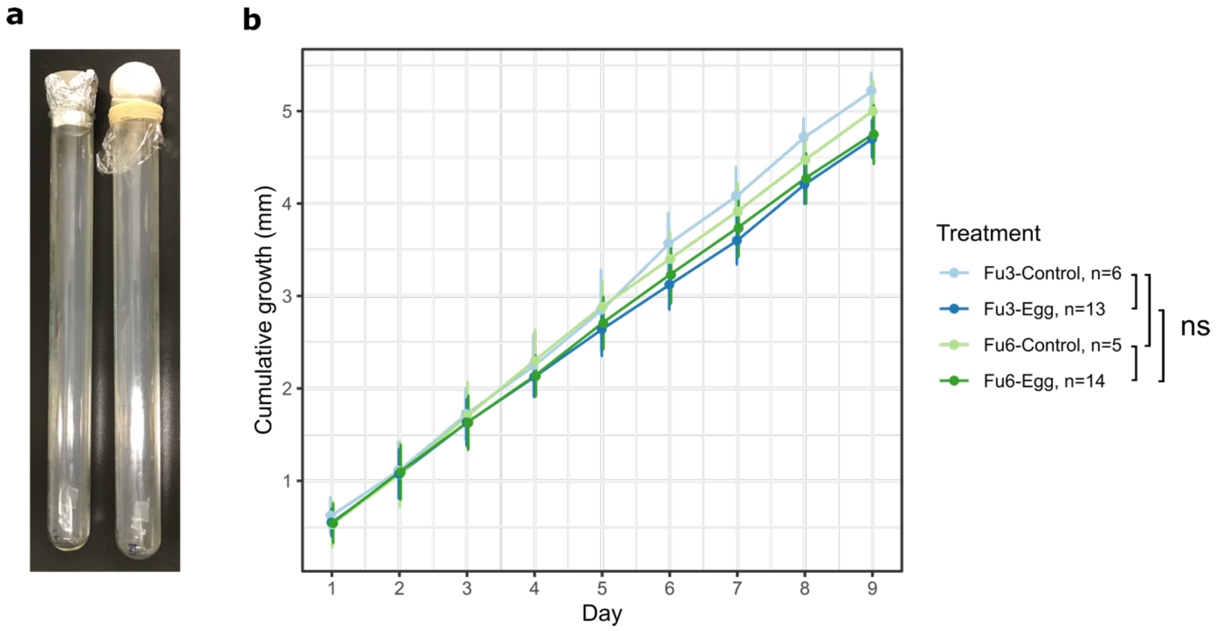

**Supplementary Figure 3.** Chinese soft-shelled turtle *P. sinensis* egg inoculated with conidia of *F. falciforme* Fu3 or *F. keratoplasticum* Fu6. (a) Fu6, four-dpi. (b) Fu3, three-dpi. (c) Fu6, three-dpi. Arrowhead indicates fungal colonization on (a) egg membrane as mycelium mass and (b and c) on the egg content as white blotches. Scale bar is approximately 5 mm.

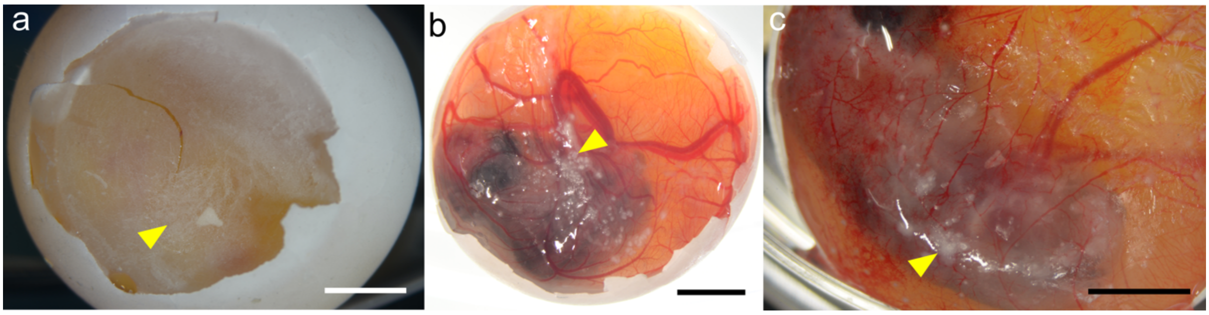

**Supplementary Figure 4.** Proportions of (a) genome features and (b) repeat contents in the six sequenced FSSC assemblies. The repeat element SINE was found in Fu3, LHS14 and Fs6 for 0.12, 0.05 and 0.05% respectively and were not noticeable in the plot.

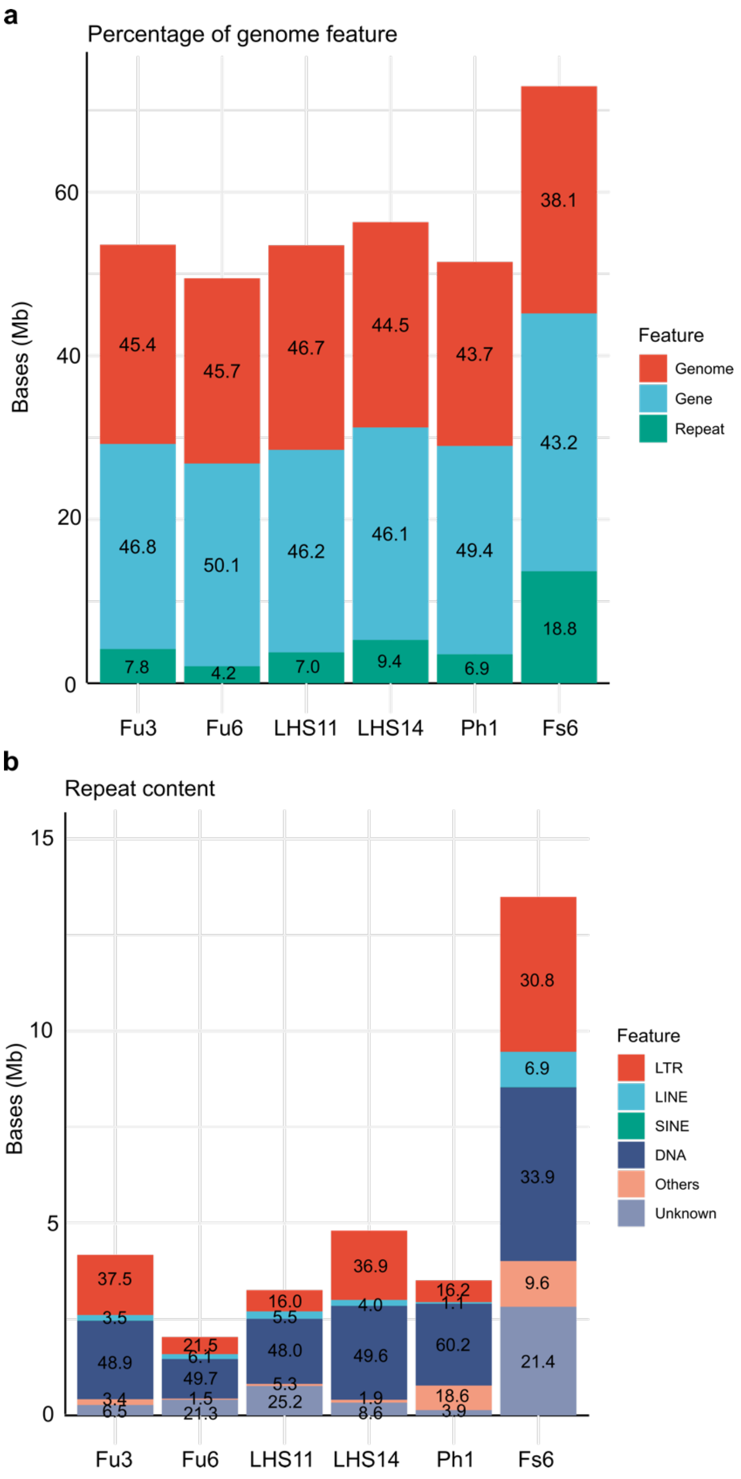

**Supplementary Figure 5.** Multi-locus sequence typing (MLST) phylogeny tree of ITS, RPB2, and TEF1 regions in FSSC. Asterisk indicates > 80% bootstraps value. Species name in bold indicates isolate sequenced in current study.

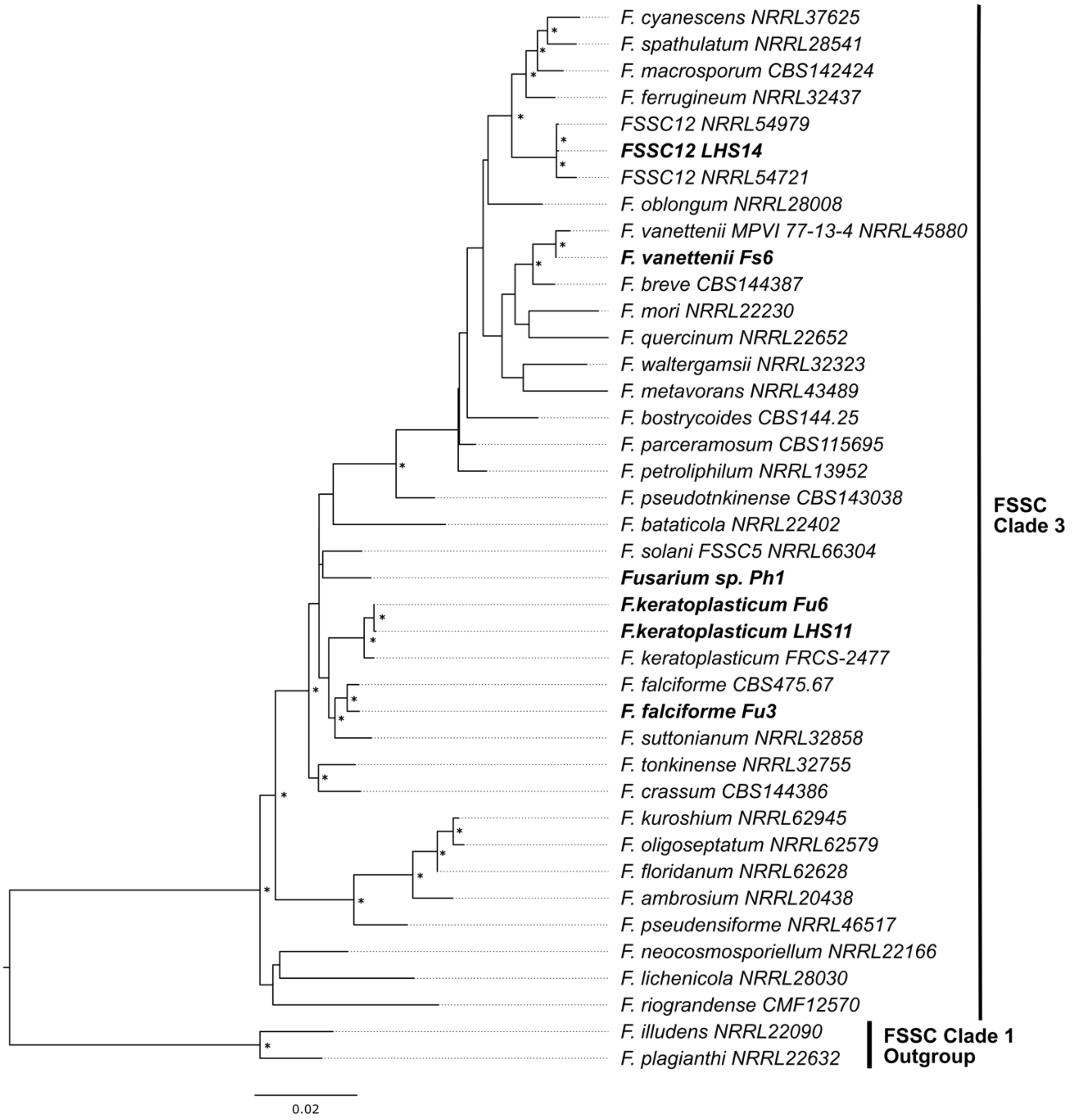

**Supplementary Figure 6.** Genome phylogeny of 23 *Fusarium* assemblies and *Beauveria bassiana* as the outgroup constructed using 2,385 single-copy orthogroup sequences. Asterisk indicates 100% bootstrap value. Coalescent unit on scale bar applies on internal branch only. Orange box denotes the FSSC clade. Species name in bold represents genomes sequenced in current study. Source origin (host) of isolates represented by icons.

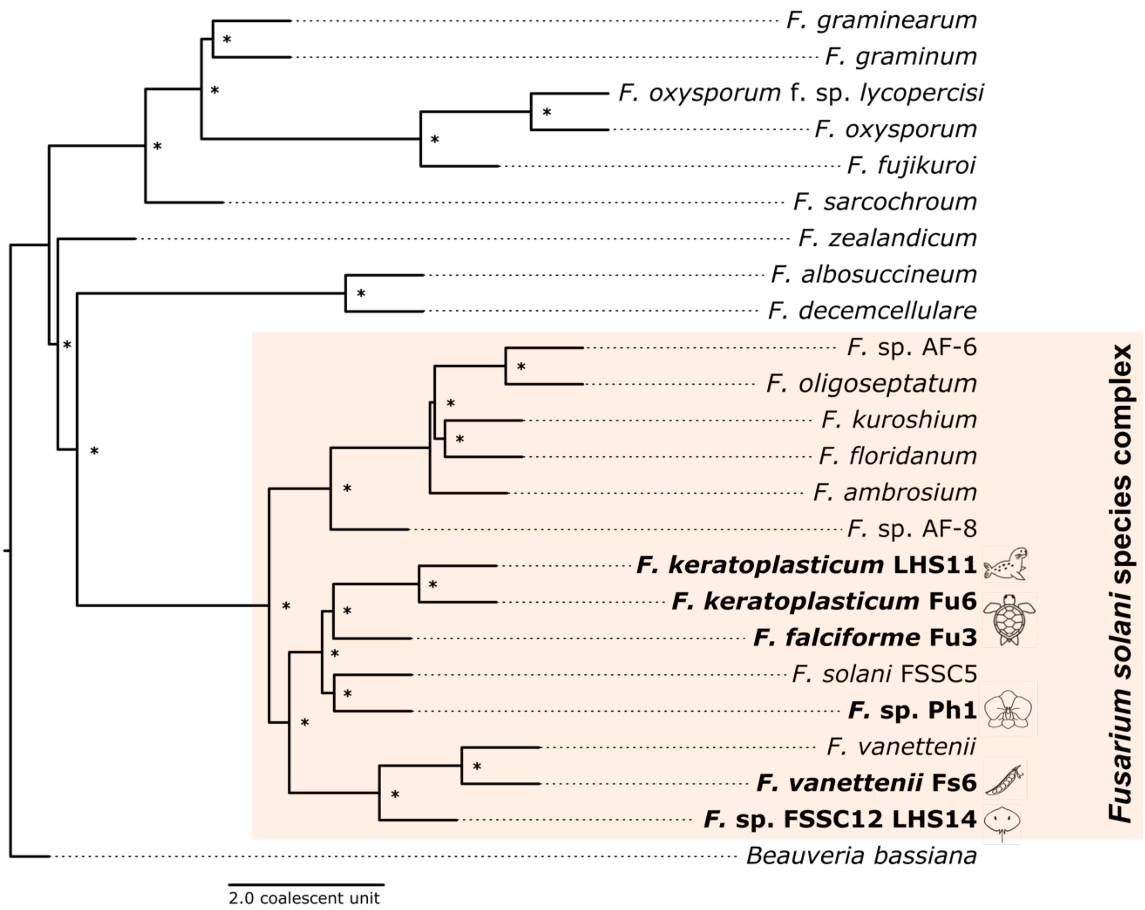

**Supplementary Figure 7.** Synteny analyses of FSSC genomes with *F. falciforme* Fu3 as reference show (a) number of single-copy orthogroup shared between chromosomes and (b) chromosomal location of these genes.

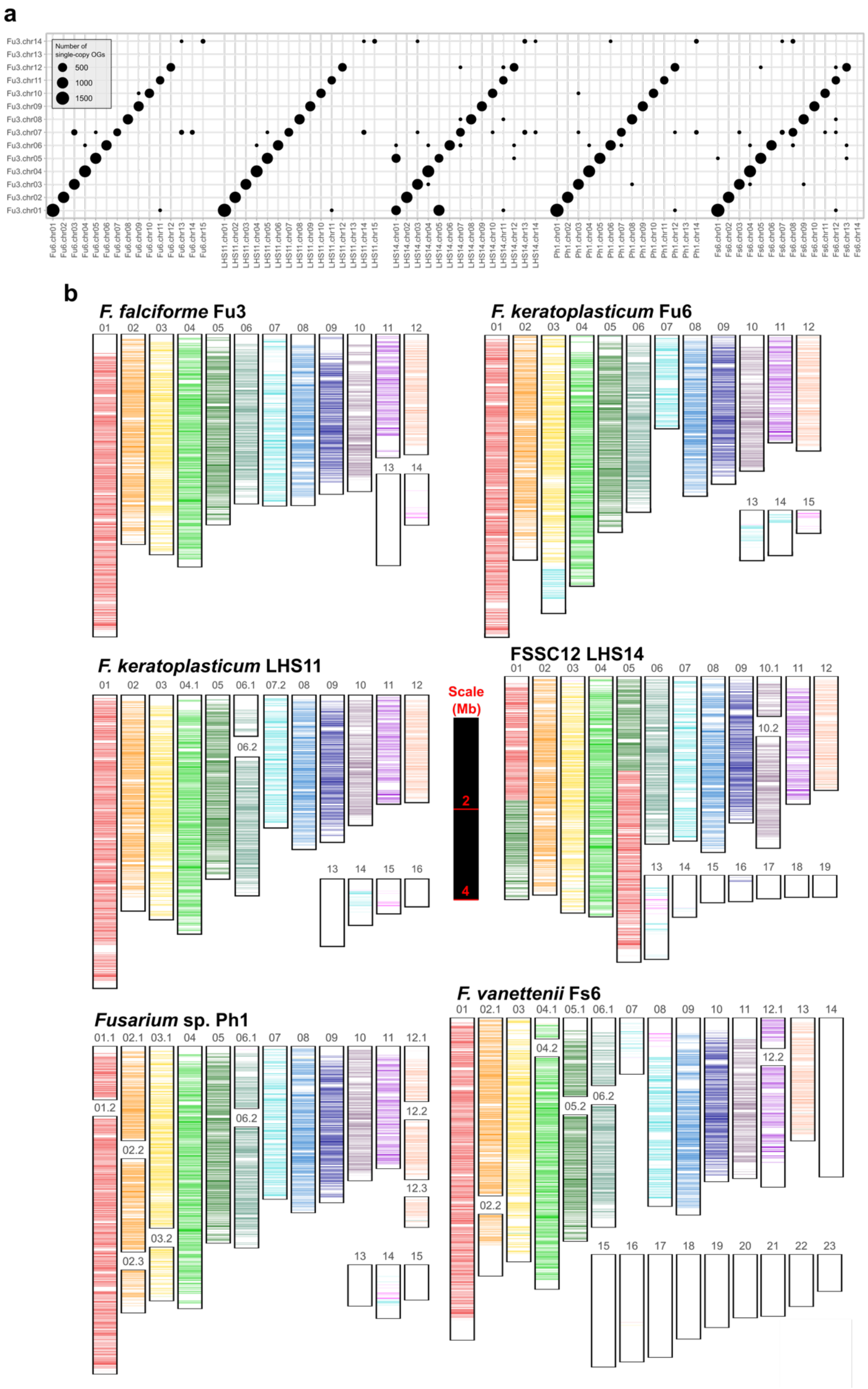

**Supplementary Figure 8.** Location of FSSC-specific genes across genomes. Single red line indicates one gene.

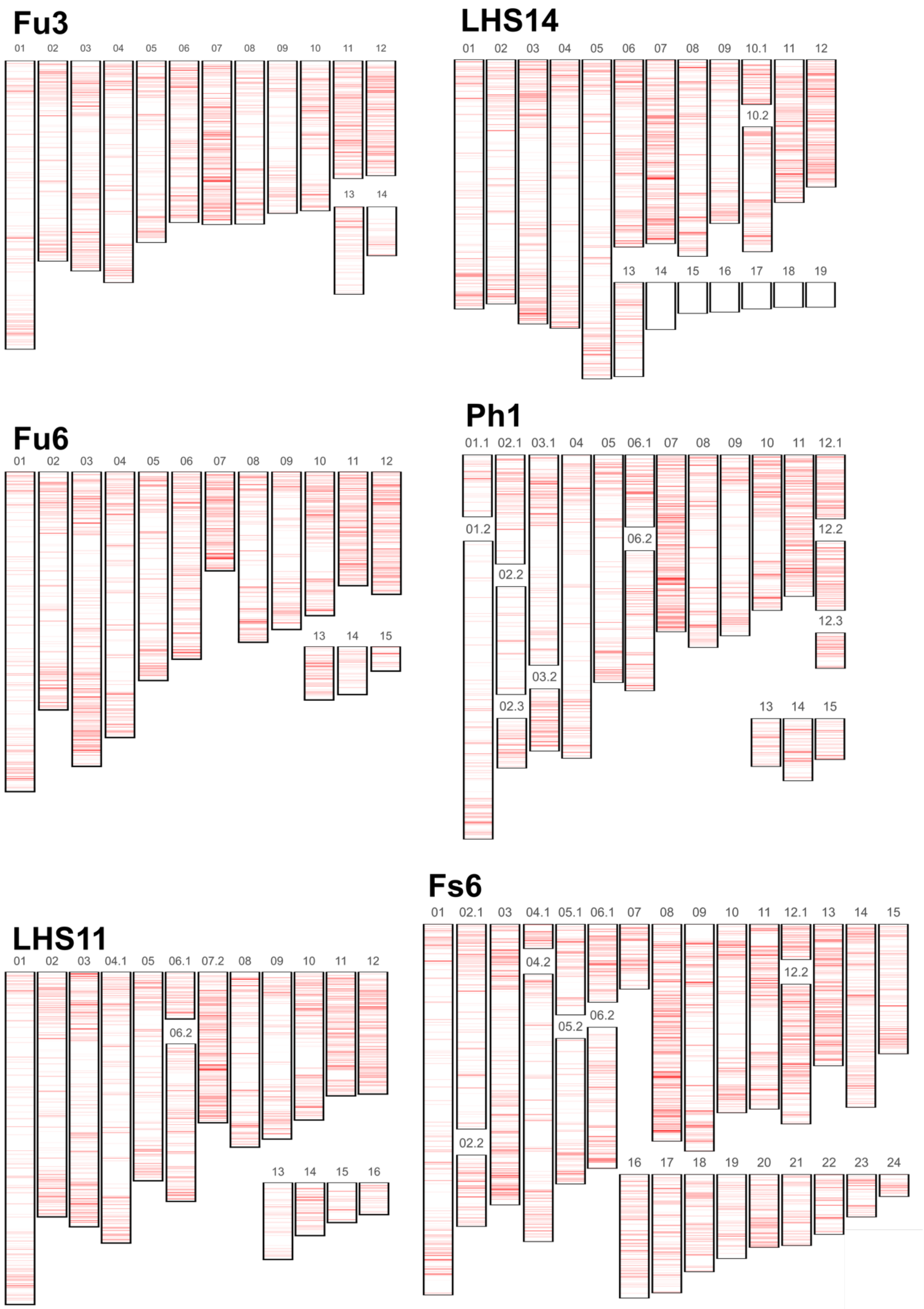

**Supplementary Figure 9.** Methylation level in (a) *F. keratoplasticum* LHS11, (b) *Fusarium* sp. Ph1, and (c) *F. vanettenii* Fs6 genomes. Plots on the left demonstrate methylation level of each chromosome and plots on the right show boxplots of each chromosome type tested for significant differences using Wilcoxon-test (****: $p < 0.0001$; ns = $p > 0.05$).

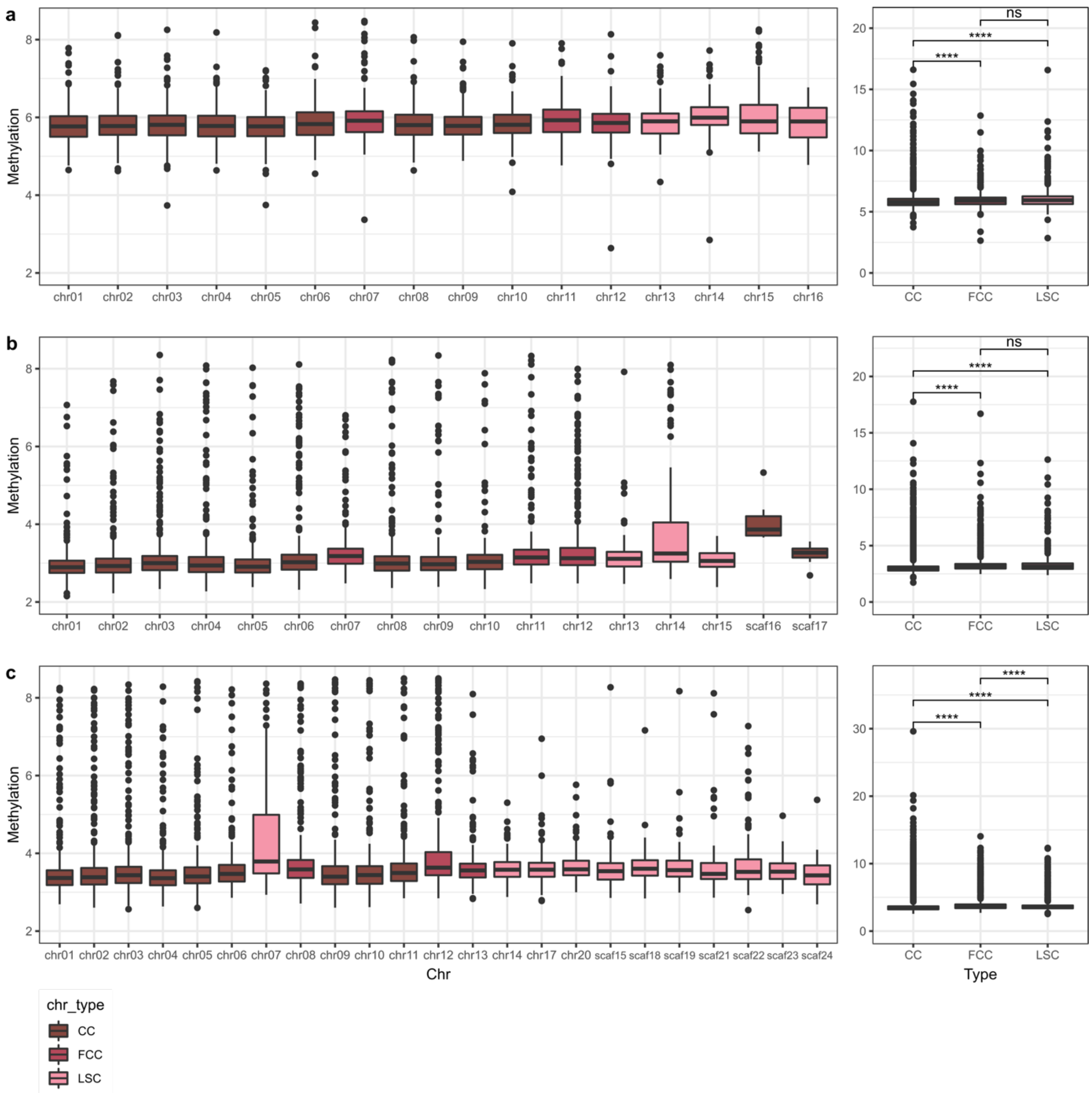

**Supplementary Figure 10.** Selection level estimates of non-synonymous substitutions (d*N*), synonymous substitutions (d*S*) per gene, and their ratio (d*N*/d*S*) comparing each group differentiating location of single-copy orthogroup between *F. falciforme* Fu3 and *F. keratoplasticum* Fu6. Groups compared include gene located on the same corresponding core chromosome ‘CC_same’ and fast-core chromosome ‘FCC_same’ while ‘Rearranged’ indicates genes not located on the corresponding chromosomes.

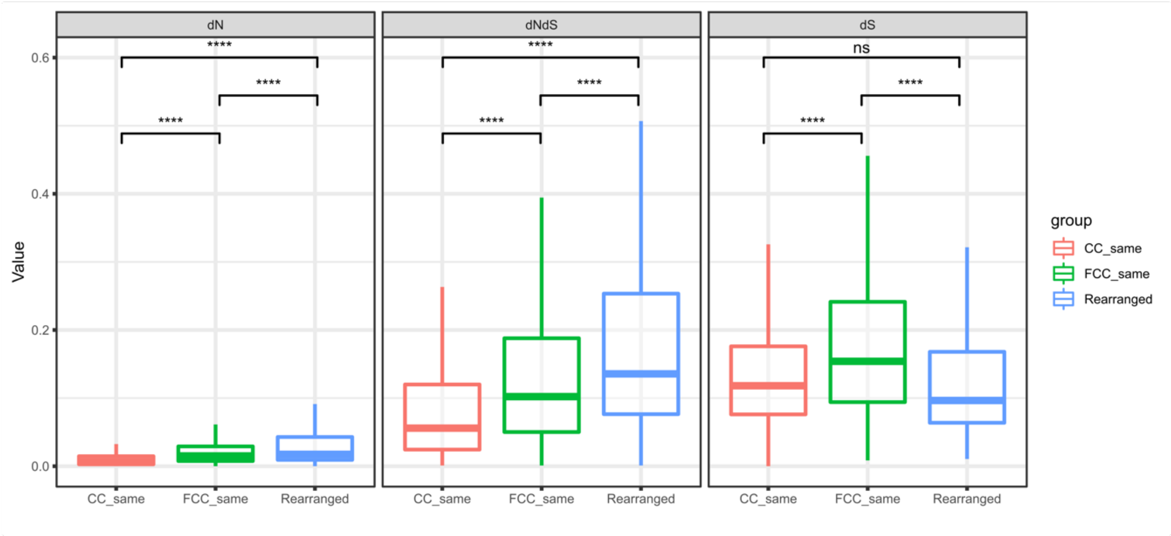

**Supplementary Figure 11.** Syntenic dotplot produced via PROmer comparing between *F. falciforme* Fu3 and (a) *F. graminearum* and (b) *F. fujikuroi*.

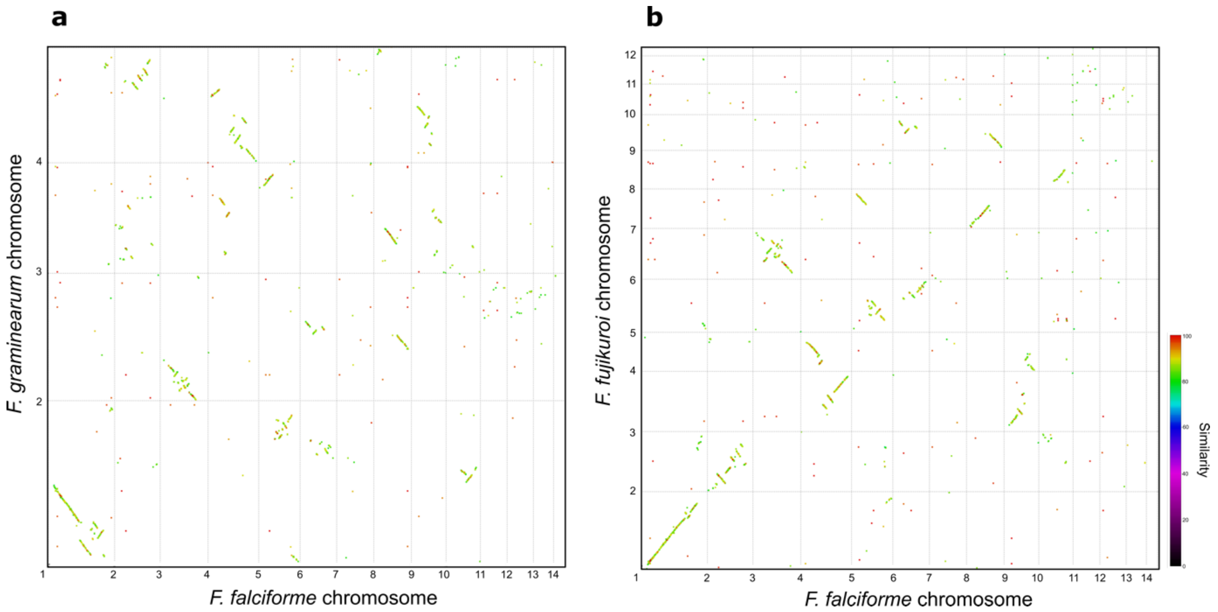

**Supplementary Figure 12.** Pairwise chromosome comparisons of shared orthogroup number between *F. falciforme* Fu3 and *F. oxysporum* f. sp. *lycopercisi* 4287.

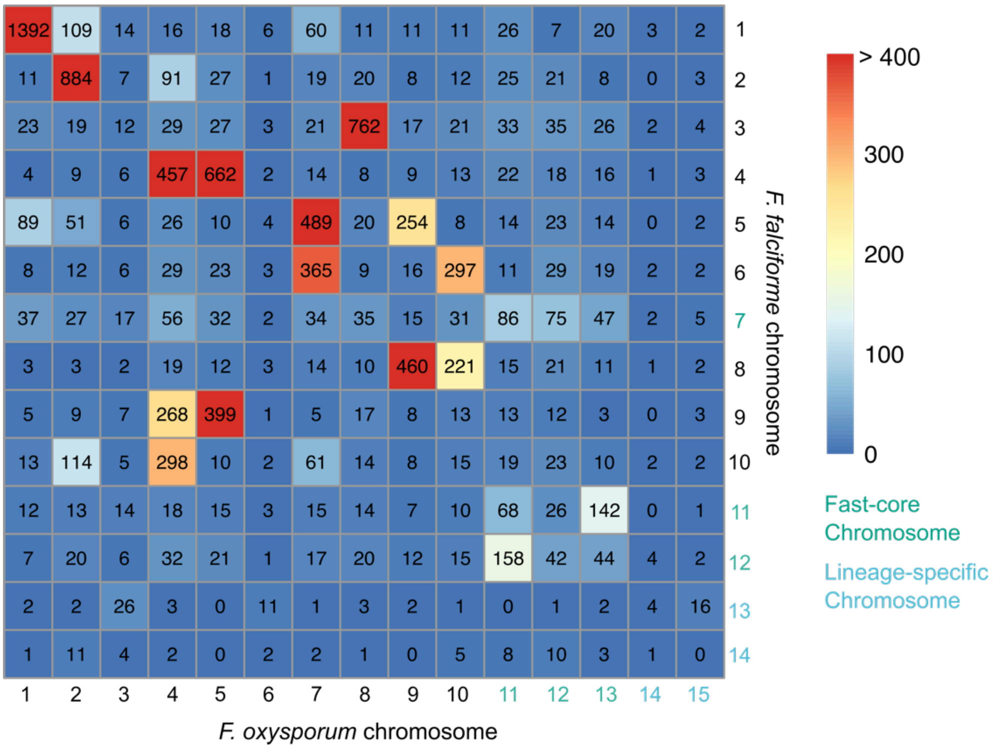

**Supplementary Figure 13.** Principal component analyses (PCA) of pathogens' gene expression pattern of all samples in (a) *F. falciforme* Fu3 (F) and (b) *F. keratoplasticum* Fu6 (K) infecting egg. 'B' and 'M' denote blotch and membrane of inoculated samples, respectively.

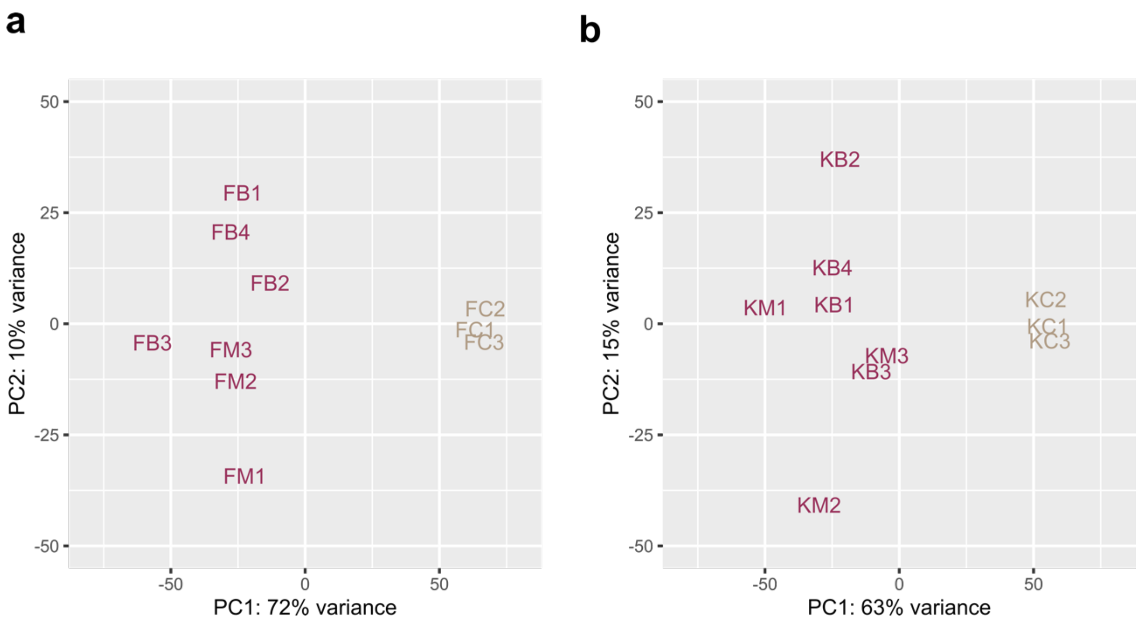

**Supplementary Figure 14.** Correlation plots of $log_2$ normalized transcript per million (TPM) of one-to-one orthologous gene of *F. falciforme* Fu3 and *F. keratoplasticum* Fu6 in (a) inoculated and (b) control (mycelium) samples.

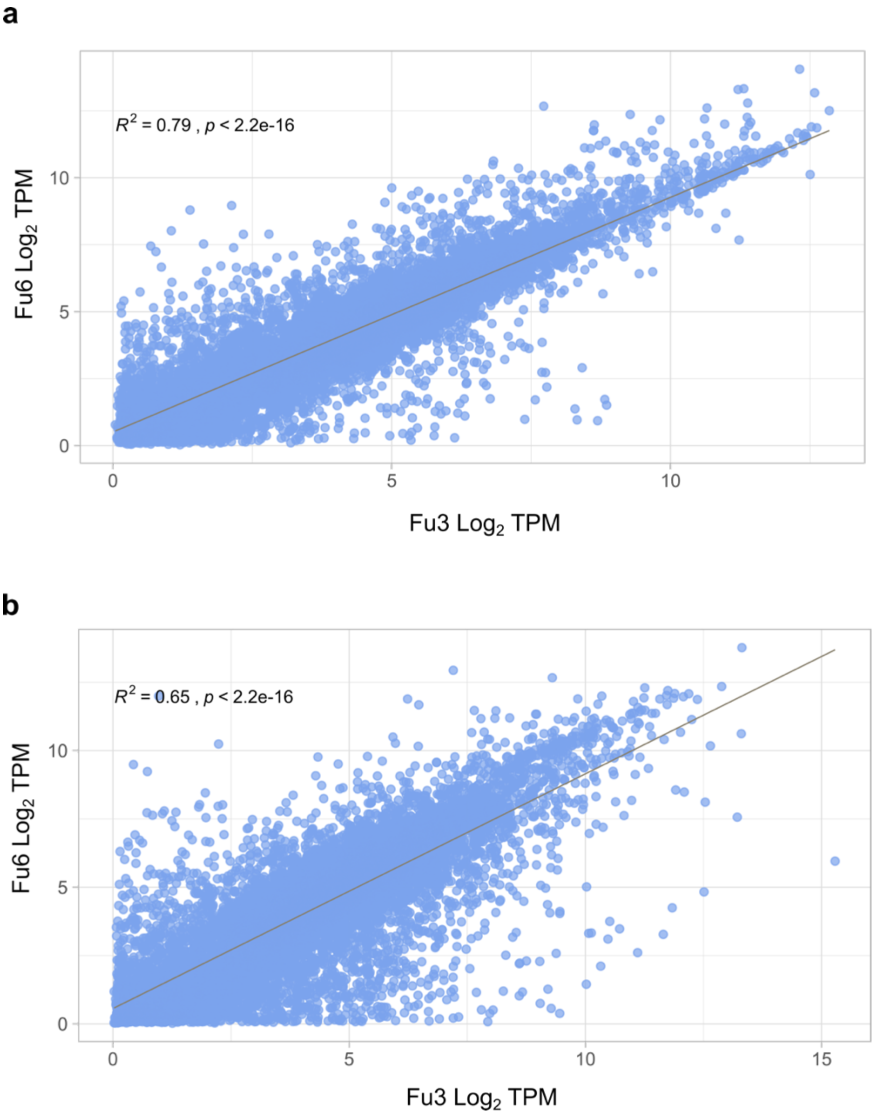

**Supplementary Figure 15.** The distribution of all differentially expressed genes of FSSC pathogens during egg inoculation experiment in (a) *F. falciforme* Fu3 and (b) *F. keratoplasticum* Fu6. Each red line represents single gene.

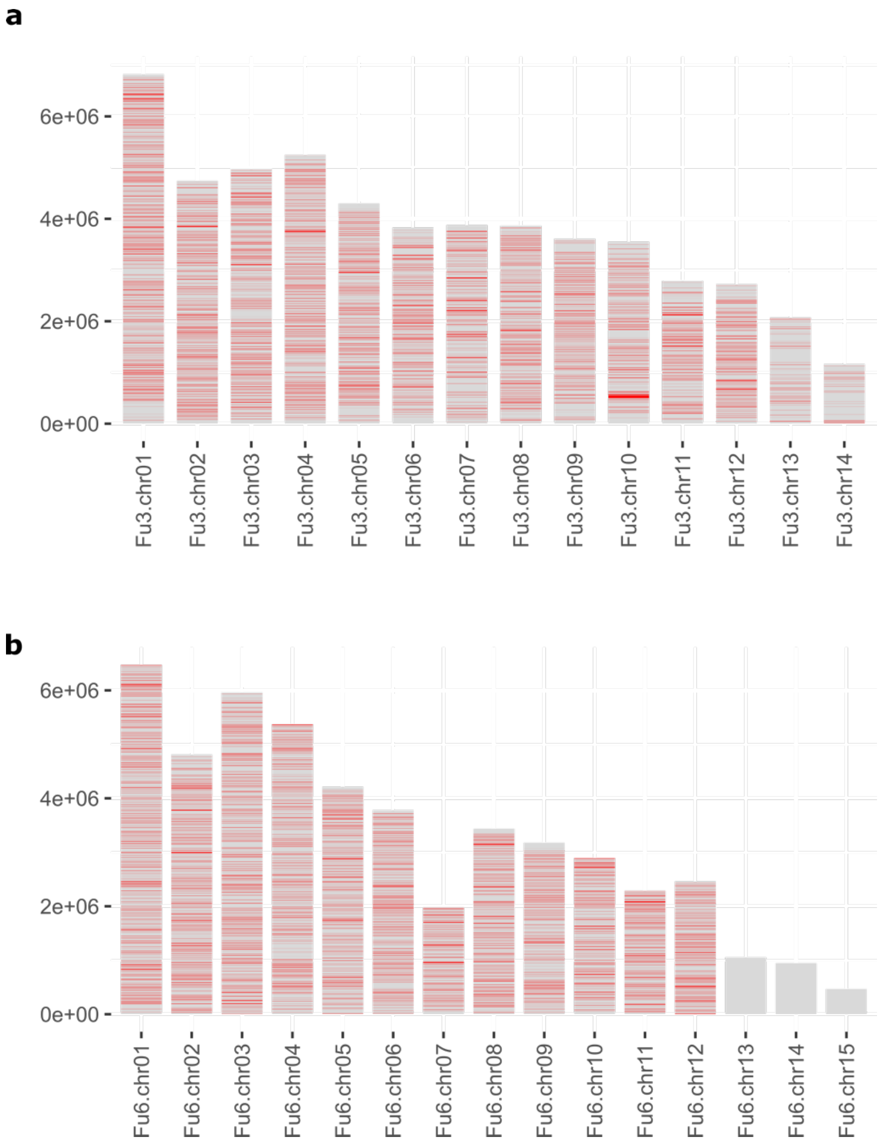

**Supplementary Figure 16.** Gene expression pattern of the animal host *P. sinensis*. (a) Principal component analyses (PCA) of gene expression pattern compared between host inoculated by *F. falciforme* Fu3 and *F. keratoplasticum* Fu6 (sample name starts with F and K, respectively) and natural developing host (DRR; Wang et al., 2013). (b) Correlation plot of $log_2$ normalized transcript per million (TPM) of host gene inoculated by *F. falciforme* Fu3 and *F. keratoplasticum* Fu6.

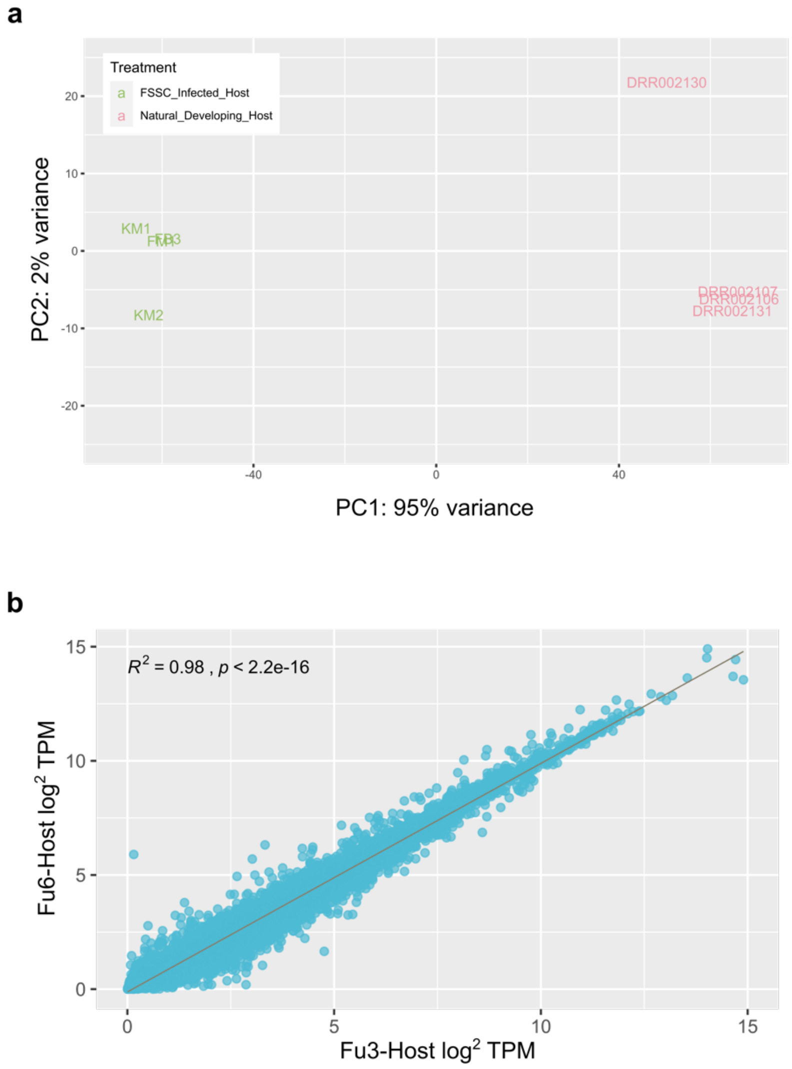
